# Modulation of SAL retrograde signaling promotes yield and water productivity responses in dynamic field environments

**DOI:** 10.1101/2025.03.10.642515

**Authors:** Andrew F. Bowerman, Marten Moore, Arun Yadav, Jing Zhang, Matthew D. Mortimer, Zuzana Plšková, Estee E. Tee, Eng Kee Au, Derek P. Collinge, Gonzalo M. Estavillo, Crispin A. Howitt, Kai X. Chan, Greg J. Rebetzke, Barry J. Pogson

## Abstract

- Chloroplast-to-nucleus retrograde signaling enables rapid stress responses in plants, but whether these signals accumulate to affect crop performance across entire growing seasons under field conditions remains unknown.
- We generated wheat mutants with targeted deletions in specific SAL gene copies from two distinct homeologous groups (TaSAL1 and TaSAL2), creating lines with enhanced stress signal responsiveness. We tested these lines across 15 field trials spanning diverse Australian environments with varying temperatures, rainfall, and irrigation regimes, measuring physiological responses, yield, biomass, and water productivity.
- Lines with TaSAL2 gene deletions showed 4-8% yield improvements with enhanced water productivity, while TaSAL1 deletions reduced yields. The TaSAL2 mutants maintained superior photosynthetic function under drought stress, showed improved relative water content, and demonstrated enhanced yield stability across environments. Canopy temperature measurements revealed dynamic stomatal regulation, with increased closure during midday stress periods but normal aperture under benign conditions. Significantly, specific SAL modifications enhanced photosynthetic efficiency and stress resilience without traditional yield penalties.
- Targeted modification of specific SAL homeologous groups can simultaneously improve both yield and stress tolerance in wheat. This demonstrates that retrograde signaling integrates environmental information across the plant lifecycle, and highlights the importance of locus-specific targeting and multi-environment field validation for crop modifications.

## Introduction

Climate variability poses an escalating threat to global food security, demanding urgent advances in crop stress tolerance (Bowerman *et al*., 2023). Modern crops face an unprecedented combination of abiotic stresses - from extreme temperatures and water availability to excess light and nutrient imbalances - often occurring simultaneously or in rapid succession. Plants coordinate their responses to these challenges through intricate cellular signaling networks, where even subtle changes in key pathways can trigger extensive transcriptional and physiological adaptations. Among these networks, chloroplast-to-nucleus retrograde signaling plays a particularly crucial role in integrating environmental cues with cellular responses (for review, see Pogson *et al*. (2008; Chan *et al*., 2016b; Crawford *et al*., 2018; Calderon & Strand, 2021)). These signals are essential for coordinating cellular responses to transient stress, and for optimal plastid operation given the number of plastid-located protein complexes that are nuclear encoded and subject to damage by oxidative stress. (Jan *et al*., 2022).

The SAL enzyme, an adenosine phosphatase (EC:3.1.3.7), regulates the levels of 3’-phosphoadenosine 5’-phosphate (PAP), a chloroplast-to-nucleus signal molecule (for review, see Jan *et al*. (2022)). SAL converts PAP into adenosine monophosphate (AMP), and is a negative regulator of excess light and drought tolerance in *Arabidopsis* (*Arabidopsis thaliana* (L.) Heynh.). Under drought or light stress, AtSAL1 becomes inactivated by oxidation, allowing PAP to accumulate and move between organelles, increasing up to 30-fold in *Arabidopsis* (Estavillo *et al*., 2011; Chan *et al*., 2016a).

Elevated PAP affects the expression of a raft of genes, partly through inhibition of exoribonucleases (XRNs) and resulting promotion of RNA polymerase II read-through at transcription termination sites (Estavillo *et al*., 2011; Gigolashvili *et al*., 2012; Crisp *et al*., 2018; Ashykhmina *et al*., 2019).

Studies of retrograde signaling in general (and SAL in particular) have largely focused on gene knockout lines in model species. Without a functional SAL protein, such knockouts constitutively accumulate PAP to high levels without stress induction, approximately 20-fold in *Arabidopsis alx8* mutants (Estavillo *et al*., 2011). These lines exhibit increased drought stress tolerance (40-50% longer survival compared to wild type) and improved water use efficiency with a decreased growth rate, dwarf phenotype, and leaf morphology changes (Rossel *et al*., 2006; Wilson *et al*., 2009). PAP itself has been implicated directly in the control of stomatal aperture closure across diverse taxa with functioning stomata, from mosses and ferns to monocots and dicots (Zhao *et al*., 2019). PAP bypasses and augments canonical ABA signaling; this includes exogenous and endogenous addition of PAP to ABA-insensitive *Arabidopsis* lines restoring stomatal closure (Pornsiriwong *et al*., 2017). Broader effects of knocking out AtSAL1 include changes in hormone levels, induction of phosphate starvation responsive gene expression, and modified metabolome including altered levels of osmoprotective compounds (Rossel *et al*., 2006; Wilson *et al*., 2009; Hirsch *et al*., 2011). Thus, determining the transient stress-responsive functions of PAP *bona fide* is confounded by the persistently elevated concentrations of PAP in mutants, which result in developmental anomalies (Phua *et al*., 2018a).

Recent studies in wheat (*Triticum aestivum* L.), which possesses a more complex SAL gene family than *Arabidopsis*, show varied outcomes from attempts to knock down all SAL copies simultaneously, including purported yield penalties alongside improved resilience. VIGS-mediated silencing improved drought tolerance markers (Manmathan *et al*., 2013), while CRISPR-based knockouts reported enhanced stress resistance based on osmotic shock experiments (Abdallah *et al*., 2022) and yield penalties in pots subject to competitive glasshouse conditions (Mohr *et al*., 2022); noting that pot-based yields are perilous measures (Passioura, 2006). None of these studies considered the roles of different SAL homeoforms, assayed any traits in the field, or measured the impacts of the genetic modification on the enzyme or its substrate. Nor did they consider whether retrograde signaling is a transient process or is integrated across the life cycle.

Indeed, more broadly retrograde signaling research has focused on standardized conditions with single stresses, rather than the complex, unpredictable stress combinations crops face in the field, and ignored if it has a cumulative effect on the entire life cycle of plants in response to multifactorial stresses (for review of multifactorial stress, see Zandalinas & Mittler (2022)). The SAL-PAP pathway’s role in coordinating chloroplast stress response, regulating hormonal signaling, and metabolic responses makes it ideal for studying these complex conditions (Phua *et al*., 2018b). However, this necessitates a different strategy to simple loss-of-function genetic modifications and single stresses in controlled environments.

The challenges we meet in this report are to first test the null hypothesis that short-term changes mediated by retrograde signaling pathways have no ongoing impact on crops grown in dynamic and varied environments. Second, to consider how do we best translate findings in a model plant under controlled environments into crops that are selected for high and stable yields across multiple environments? Our strategy was to create knockouts of select TaSAL homeologues in wheat. SAL is typically in excess (Chan *et al*., 2016a), which means that PAP levels remain low unless the stress is significant, or AtSAL1 transcript levels are reduced by more than 80% (Phua *et al*., 2018a). By reducing one to two of the six homeologues present in cv. Chara, we engineered plants with slightly diminished overall SAL activity, thereby lowering the threshold for PAP signaling. We postulate that such primed plants would respond more dynamically to environmental perturbations. We utilized heavy ion bombardment (HIB) to knockout *TaSAL* genes, enabling us to assess these materials in natural environments during a period when gene-edited field trials were not authorized (2015-2019) and producing a crop that is globally far more deployable to industry. Our evaluation of the primed retrograde signaling plants in natural environments was conducted across 15 distinct settings, covering four different cereal-growing districts over five years, characterized by varying rainfall and irrigation regimes. Subsequently, we incorporated the observations into a multi-environment trial (MET) model to examine whether modulation of retrograde signaling influences altered lifetime responses to diverse and dynamic natural environments. An additional aspect of the study involves an evaluation of the trade-offs in yield resilience for genes that can modulate both stress responses and developmental processes.

## Materials and Methods

### Phylogeny and expression analysis

Sequence homologues of AtSAL1 (NP) were identified from Liliopsida using Uniprot BLAST (e-value<0.1). A sequence similarity network was constructed and filtered to retain sequences with: 2 40% identity that clustered with *Arabidopsis* (*Arabidopsis thaliana* (L.) Heynh.) SAL1/2/3/4 references, and sequences between 250-600 amino acids in length. Wheat (*Triticum aestivum* L.) sequences were updated using homologues identified from Gramene, and AHL enzyme sequences were added as an out-group. Multiple sequence alignment was performed using T-COFFEE, refined with MAFFT, and manually curated. Sequences with non-consensus insertions/deletions were removed. Phylogenetic reconstruction used ModelFinder for model selection, with multiple replicates and ultrafast bootstraps. The final phylogeny was selected based on outgroup contiguity and visualized in R. Detailed protocols and software versions are provided in Supporting Methods S1.

Expression data were retrieved from the Wheat RNA-seq Database https://plantrnadb.com/wheatrna/ (Zhang *et al*., 2020) and from the database of Loudya *et al*. (2021). Gene co-expression analysis was performed using ATTED-II, using all members of each TaSAL group as seed sequences. The top 100 co-expressed genes for TaSAL1 and TaSAL2 were then used in ShinyGo v0.80 to summarise GO terms related to these co-expressed genes (Ge *et al*., 2020).

### Identification of *TaSAL* mutants and generation of double null lines

A wheat population mutagenised by heavy ion bombardment (HIB) was used to search for mutations in each *TaSAL* homeologues (Regina *et al*., 2015). Rather than single nucleotide changes which are seen in chemical mutation-based populations, HIB mutations result in large deletion mutations, usually encompassing entire genes. A total of 3000 individual M3 HIB lines were screened using homeoform specific primers for deletions in each *TaSAL* gene by PCR, resulting in the identification of five primary null lines, three deletions on chromosome 4A and two on chromosome 5D. Primer sequences are listed in Supporting Information Table S1. Multiple crosses were generated using combinations of each of these null lines; all line details are found in Supporting Information Table S2. Deletions were characterised using two targeted genotype-by-sequencing (tGBS) approaches, completed by Agriculture Victoria (AgriBio, Bundoora, Australia). Briefly, one tGBS assay was constructed from a subset of the Infinium 90K SNP chip (Chidzanga *et al*., 2021), and the second tGBS method was constructed using exome sequencing results from approx. 1000 wheat lines (Sharma *et al*., 2022). Combined, this maximised the marker density and maximised our chances of identifying deletions. See Supporting Information files details of deletion characterisation.

### Plant growth and drought stress in glasshouse

All lines were grown in a controlled greenhouse with day-length extension to 16h at 24/16°C day/night temperature and 60% relative humidity. Pots of 25 cm (diameter) × 30 cm (h) were filled with compost-based potting mix and arranged in a randomised complete block design. All pots were watered daily to pot capacity until drought stress was imposed at the 3-4 leaf stage (21 days). Two drought regimes were used. **Cyclic Drought**: At day 0 of drought, water was withheld in droughted plants until day 21 of the first drought and then re-watered to their 100% soil water capacity as per Bowne *et al*. (2012). **Terminal drought**: All pots were saturated with water at day 0 and designated drought-stressed plants were not watered up to 42 days of drought. All control plants were maintained at 100% water capacity. Where required, soil weight measurements were collected to compare water use during drought periods.

### Field trial management

Performance of genotypes was assessed under a series of field trials undertaken across four seasons at four managed environment facilities in New South Wales and Victoria, Australia. Details of facilities can be found in Rebetzke *et al*. (2013), and are indicated, along with temperature and rainfall variation between and within sites in Figure 4. Multiple trials were performed in each site/year with varying access to irrigation to ensure variation in water availability between trials. All designs were produced using a complete randomised block design approach. In all experiments, all the plots were treated with 30 mm of pre-sowing irrigation and entries were sown at an optimal 3 to 5 cm sowing depth into 6-m long, 0.17-m spaced, 8-row plots at seeding rates consistent with local practice (i.e. 120 seeds per m^2^ at Narrabri, and 180 seeds per m^2^ at Yanco and Condobolin). Where irrigation was supplied, it was calculated at the site to target 25% above mean rainfall. Nutrients were supplied at sowing as Starter 15^®^ (14% N: 12.7% P: 11% S) applied at 103 kg.ha^-1^. Additional nitrogen was applied as needed to meet crop demand. Plots were end-trimmed to 5.4 m length and border rows removed prior to machine harvesting for plot yield estimates.

### PAP quantification, SAL activity assay and protein quantification

PAP was quantified using a modified version from Ashykhmina et al. (2019). Metabolites were extracted from liquid nitrogen-frozen leaf tissue (30-50 mg) using chloroform:methanol and citrate-phosphate buffer extraction. PAP was quantified by HPLC following chloracetaldehyde derivatization using commercial PAP as a standard (Sigma-Aldrich, A5763). Separation was performed on a Kinetex XB-C18 column (Phenomenex) using tetrabutylammonium hydrogensulfate and acetonitrile buffers. SAL activity was measured using protein extracts from leaf tissue incubated with PAP substrate at 37°C, with enzymatic activity determined by measuring PAP degradation via HPLC. Detailed protocols are provided in Supporting Method S2.

### ABA quantification

Leaf ABA was quantified using a modified extraction protocol. Approximately 100 mg of flash-frozen leaf tissue was homogenized in a Tissue Lyser II (Qiagen, Germany) with a steel ball bearing at 19.5 Hz for 2 minutes. The tissue was extracted twice with 2 mL cold 80% methanol by overnight shaking (25 rpm, 4°C) followed by centrifugation (17,000 rcf, 4°C). After each extraction, supernatants were vacuum concentrated to 500 µL using a miVac Duo Concentrator (GeneVac, UK). A final methanol wash step was performed and concentrated to 500 µL. Combined concentrated extracts (1.5 mL total) were further reduced to 500 µL under vacuum. Extracts were acidified to pH 2.5 with 0.5 mM HCl and partitioned against equal volumes of ethyl acetate. The organic phase was dried under vacuum and resuspended in 1 mL Tris-Buffered Saline (TBS). ABA was quantified using the Phytodetek ABA Kit (Agdia Inc, USA) according to manufacturer’s instructions, with absorbance measured at 405 nm (Tecan Group, Switzerland). Final ABA concentrations were expressed as ng/g dry weight.

### Photosynthesis, stomatal conductance, relative water content and carbon isotope discrimination

An open gas-exchange system with a 2 × 3 cm clamp-on leaf cuvette was used to simultaneously measure stomatal conductance and photosynthesis (Li-6400; Li-Cor, Inc., Lincoln, USA). The topmost expanded leaf was chosen for taking measurements. Measurements were taken after leaves were adapted in the chamber for approximately 10 minutes at 380 µmol CO_2_ and light intensity of 1000 µmol/m/s, block temperature of 24°C, and ambient humidity approximately at 55%.

Relative water content was measured as in Wilson *et al*. (2009), with leaf sections cut and weighed to measure fresh weight (F_W_). Leaves were incubated in water at 4°C in dark conditions, blotted, and re-weighed to measure turgid weight (T_W_). Finally, leaves were dried at 80°C overnight, weighed to determine dry weight (D_W_), and relative water content was calculated as

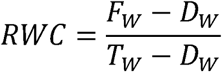

Samples of dried and ground plant material were weighed to 1.5 mg (within 0.2 mg) into folded tin foil cups. These were then combusted in a Carlo-Erba EA-1110 Elemental Analyser (Carlo Erba, Milan, Italy) providing a controlled oxidation and reduction. Gases were separated in the helium carrier using a packed gas chromatograph (HayeSep D 80/100, Grace Davison, Australia) at room temperature and passed to a stable isotope ratio mass spectrometer (Isoprime, Micromass U.K., Manchester, U.K.). A pulse of reference CO_2_ succeeded each CO_2_ sample peak, to correct for pulse to pulse variation.

Standards used were a C3 beet sucrose (Beet89) with a δ^13^C of −24.62‰ Vienna Peedee Belemnite (VPDB) and ANU C4 cane sucrose with an accepted value of −0.45‰ VPDB. Each run commenced with two of each sucrose standard and proceeded with pairs of standards every 10 samples. Final corrections were made using the slope and offset derived from the two standards. Values were then converted to Δ form using

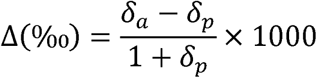

where δ_a_ and δ_s_ are the δ^13^C values for atmosphere (−8‰) and the sample, respectively (Farquhar *et al*., 1989).

### Hyperspectral imaging and leaf temperature measurements

Hyperspectral measurements were taken using ASD FieldSpec 3 (Boulder, CO, USA). Photosynthetic parameters were then estimated using the Wheat Physiology Predictor at https://wheatpredictor.appf.org.au/ (Furbank *et al*., 2021).

Canopy temperatures were measured at Narrabri, NSW, on two consecutive days in 2017 and once in 2019 using infrared cameras deployed on unmanned aerial vehicles (UAVs). Six measurements were taken over seven hours of daylight to capture diurnal temperature patterns. Data were analysed using multivariate repeated measures linear mixed effects models, with plot as a random effect and time, genotype, and water treatment as fixed effects. Best linear unbiased estimates (BLUEs) were calculated for each time point.

### Statistics

All statistics were performed using R (R Core Team, 2021). Field analysis was performed using ASREML-R (Butler, 2022) or the ‘lme4’ package (Bates *et al*., 2015). Multi-environment trials were analysed using the two-stage approach of Smith *et al*. (2001) facilitated by the use of the ‘ASRtriala’ package (Gezan *et al*., 2022). Models best fitting a trait in each trial were first constructed before combining using the Factor Analytic framework, identifying the optimal number of factors for each trait.

## Results

### Identification of null alleles in *TaSAL* subtypes

Previously 5-7 SAL genes were identified in different wheat varieties (Mohr *et al*., 2022). Phylogenetic analysis of monocot SAL homologues showed independent duplications in both Pooideae and Panicoideae subfamilies of the Poaceae, with retention suggesting functional importance of both groups (Figure 1a). We designated these as TaSAL1 (5A/B/D) and TaSAL2 (4A/7A/7D), with the 4A copy resulting from a known 4AL/7BS translocation (Devos *et al*., 1995). Chara cv., used herein, contained six copies across two paralogous groups: 5A/B/D and 4A/7A/7D.

**Figure 1.**
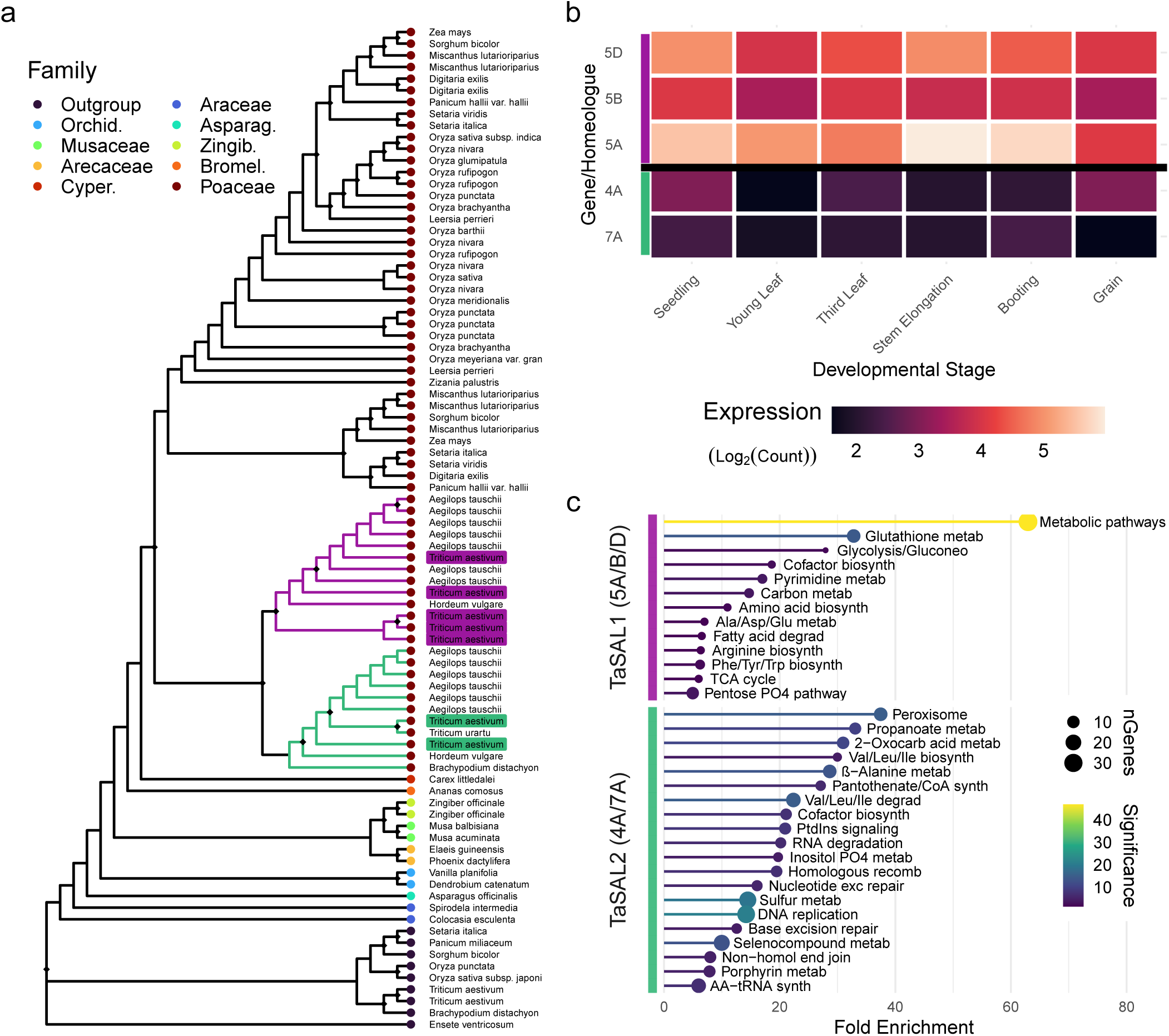
Wheat SAL gene diversity. (a) Phylogeny of Liliopsida SAL sequences. Black diamonds indicate bootstrap support >95% based on 1000 ultrafast bootstrap replicates. Purple: TaSAL1 (chr 5), green: TaSAL2 (chr 4A/7A/7D). (b) Expression of wheat SAL genes. (c) GO enrichment for TaSAL groups. Significance = -log_10_(*p*-value).

Analysis of public RNA-seq based expression data revealed that TaSAL1 homeologues showed 10-fold higher expression than TaSAL2 and different developmental profiles (Figure 1b, Supporting Information Fig. S1) (Loudya *et al*., 2021). Co-expression analysis of the TaSALs revealed enrichment in distinct functional associations: TaSAL1 with sulfur metabolism (11-fold), DNA repair (15-fold), and aminoacyl-tRNA pathways (62-fold), while TaSAL2 associated with carbon (14-fold) and amino acid metabolism (37-fold), suggesting roles in photosynthesis and energy metabolism (Figure 1c).

Given this evidence of differential expression and coexpression, we aimed to investigate the relative impact of targeted reductions in TaSAL1 and TaSAL2. We screened a Chara HIB population (M3 generation) for single gene knockouts (Supporting Information Fig. S2). Five lines were identified, two in TaSAL1-5D (designated SAL1.1 and SAL1.2) and three in TaSAL2-4A (SAL2.1, SAL2.2, and SAL2.3), and systematic crosses between these lines were generated (Supporting Information Table S2).

Genomic characterisation of background deletions in the 12 lines was performed using two targeted Genotype-by-Sequencing (tGBS) methods to genotype these lines and identify deletions. The GBSs confirmed the PCR demonstrating that all lines had complete deletions in the respective TaSAL homeologue (Supporting Information Dataset 1). The size of the deletions varied from minimal additional deletions of other loci, through to line SAL2.1, which was shown to have an additional large deletion in Chromosome 1B. Other than this line and its associated double mutants, no other incidental mutations were shown to alter traits analysed in this study, nor were any other gene deletions identified in all mutants that would be consistent with the observed phenotypes.

### Selective knockout of SAL genes lowers SAL activity under stress

To ascertain the effects of SAL mutations on PAP concentrations and enzymatic activity, lines were assessed under controlled glasshouse conditions with adequate water and drought schedules (detailed in Methods). The quantification of PAP levels in wheat has not been previously reported and required optimisation of HPLC conditions, including column selection (Kinetex XB-C18 vs standard C18), gradient simplification, and increased acetonitrile concentration to improve separation of co-eluting compounds that interfered with PAP detection.

Under sufficient water, there was no significant difference in PAP levels (Supporting Information Fig. S3) or SAL activity between mutant and control lines. However, under moderate Cyclic Drought, most mutant lines showed significant decreases in SAL activity of up to 60% compared to Chara, and several double mutant lines showed increased PAP accumulation at anthesis (Figure 2, Supporting Information Fig. S3). These genotype-dependent differences were confirmed statistically (genotype:treatment interaction: PAP content F_(12,52)_=2.42, *p*<0.05; SAL activity F_(12,52)_=4.01, *p*<0.001).

**Figure 2.**
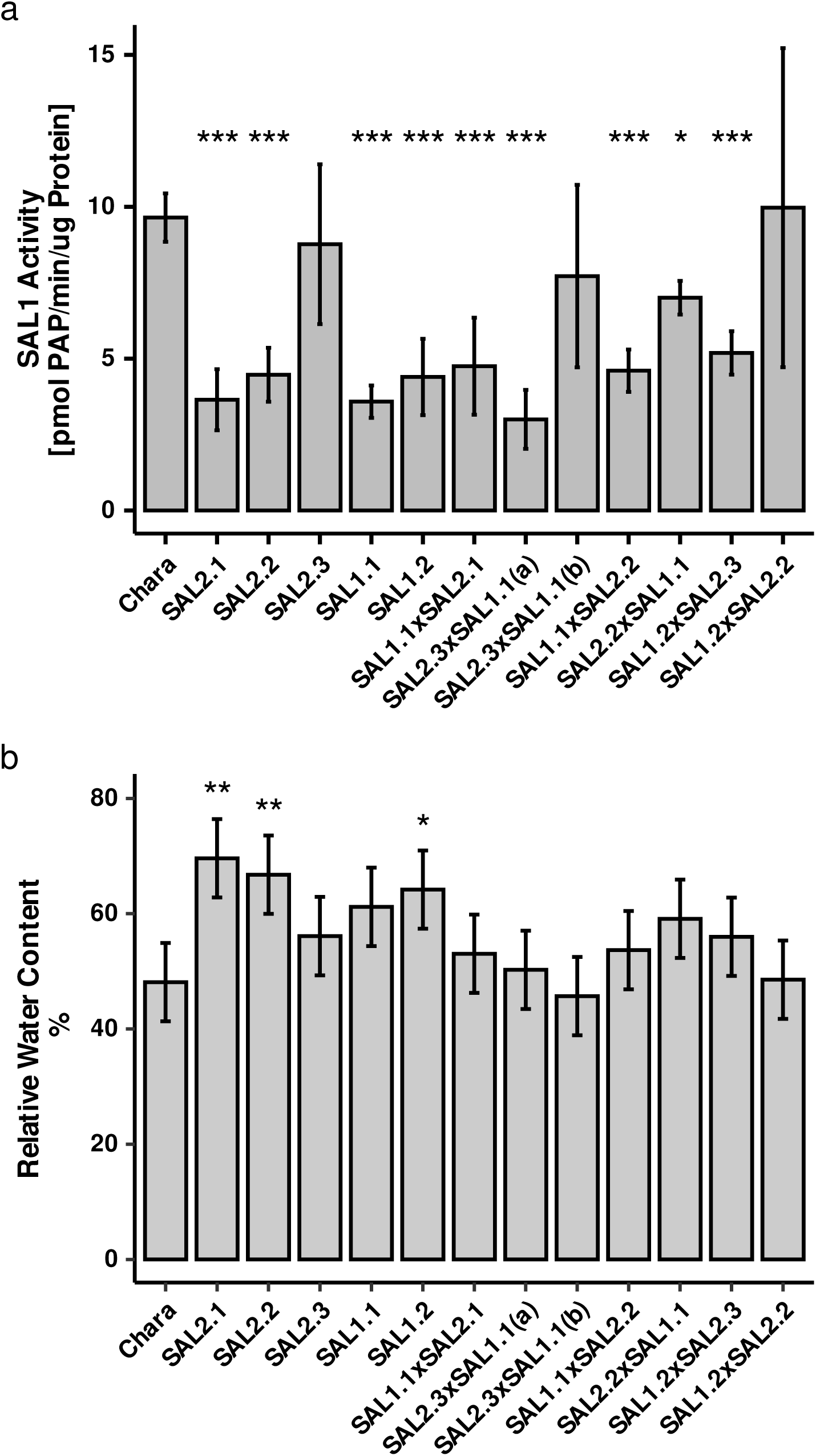
TaSAL activity (a) and relative water content (b) under drought stress. (a) SAL activity after 21d drought. Mean ± SD. Two-way ANOVA with FDR correction. (b) Relative water content after 21d drought. Mean ± Half Least Significant Difference (HLSD) from ASREML-R repeated measures model. \**p*<0.05, \*\**p*<0.01, \*\*\**p*<0.001.

Relative water content (RWC) was measured at 7, 14 and 21 days under the Cyclic Drought regime (Figure 2, Supporting Information Fig. S4B), and modeled using ASREML-R as a repeated measures experiment by using a uniform multivariate model. By day 14 four of five single nulls demonstrated increased RWC compared to Chara, as did two double null lines. At 21 days, two of three TaSAL2 lines maintained this water advantage, as did one of two TaSAL1 null lines.

To determine whether improved drought performance resulted from reduced water use or genuine stress tolerance, we analysed pot weight depletion during drought stress (days 0-25; Supporting Information Fig. S5). Mixed-effects modelling revealed no significant genotypeXtime interaction (F_(12,481)_=0.059, *p*=1.000), with water depletion rates ranging from 1.26 ± 0.23%/day to 1.46 ± 0.23%/day across all genotypes. No significant differences were detected between any TaSAL mutant and Chara.

These findings demonstrate that all genotypes experienced equivalent rates of soil water depletion, ruling out water use avoidance as an explanation for improved drought performance. The data support a tolerance mechanism whereby TaSAL mutants maintain superior physiological function whilst experiencing identical soil water. This priming of the SAL-PAP pathway creates lines that respond more dynamically to drought while maintaining normal function under unstressed conditions.

### Physiological analysis of unstressed and stressed TaSAL germplasm

No obvious changes to crop phenological traits were identified in TaSAL germplasm. Single mutants maintained comparable biomass, plant height, and tillering. From the 3rd leaf stage, plants were grown in two different watering regimes for 35 days. The well-watered regime was to sufficiency (normalized to pot weight), while drought was enforced by the Cyclic Drought regime. Single mutant lines had comparable plant height to Chara within regimes, whereas most double mutants were reduced in stature (Supporting Information Fig. S4A). The Treatment and Genotype interaction was significant (F(_12,43.4_) = 2.6, *p <* 0.05), indicating that drought stress had differential impacts on genotypes, with TaSAL mutants maintaining higher relative water content than Chara under water limitation (Supporting Information Fig. S4A).

To assess drought effects on photosynthesis (A) and stomatal conductance (g_s_), we subjected four lines at the 5/6 leaf stage plants to two regimes, well-watered (control) and withholding all water for up to 42 days (Terminal Drought regime). While photosynthesis remained stable in the control regime (Figure 3, Supporting Information Fig. S6), it declined significantly in Chara after 28 days of drought. TaSAL deletion lines maintained higher photosynthetic rates compared to Chara over the drought period (F_(3,11)_ = 6.67, *p =* 0.008). Stomatal conductance was also examined in the same experiment. While drought reduced g_s_ across all lines (F_(1,24)_ = 89.25, *p <* 0.0001, 7^2^= 0.53), there were no genotype-specific differences. This maintenance of photosynthesis despite similar g_s_ reductions suggests improved photosynthetic capacity under water limitation in TaSAL mutants as seen in Figure 3.

**Figure 3.**
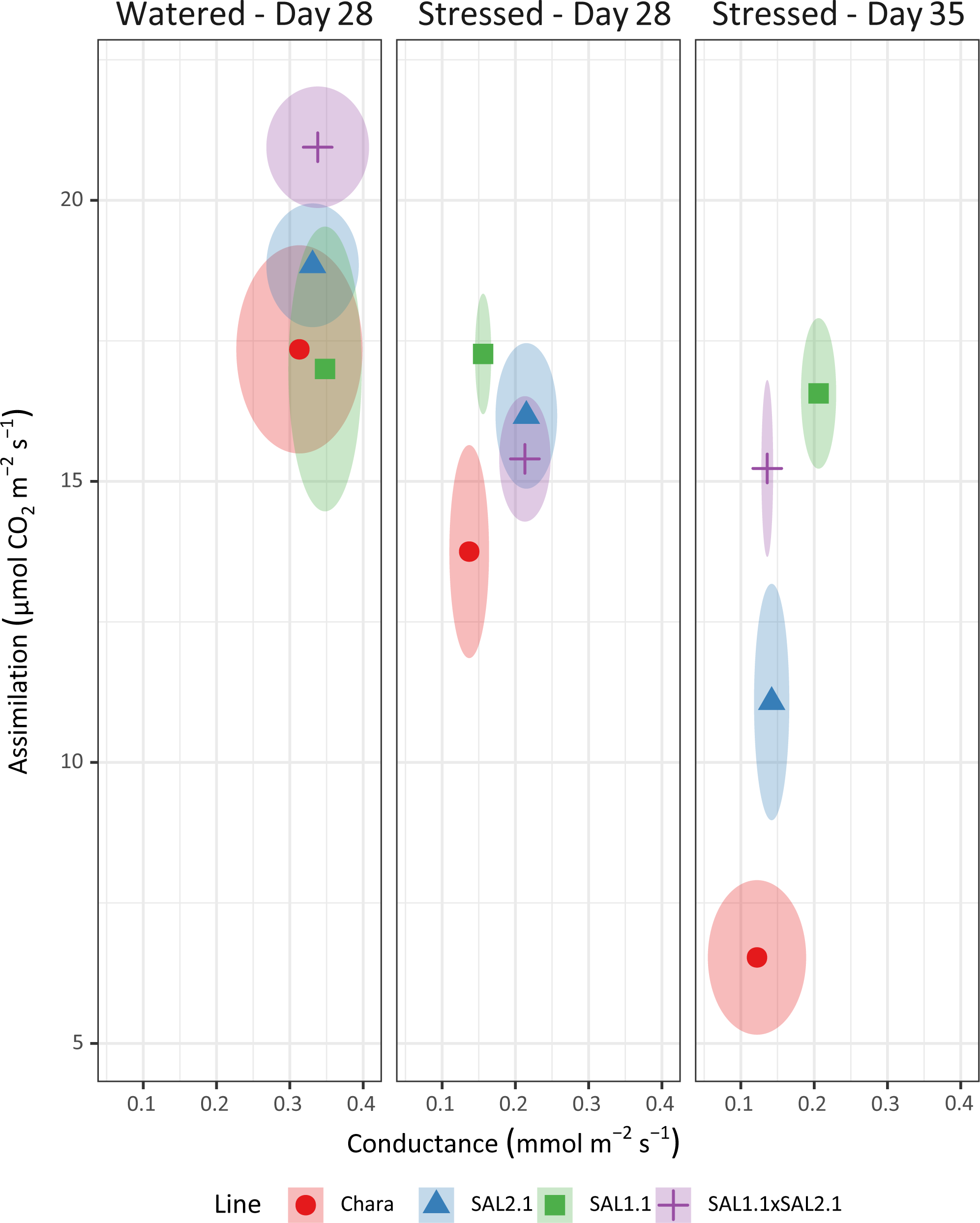
Photosynthesis vs stomatal conductance under water stress. Relationship between photosynthetic assimilation and stomatal conductance under watered control (Day 28) and Terminal Drought conditions (Days 28, 35). TaSAL mutants maintain higher assimilation rates despite similar conductance reductions compared to Chara. Mixed effects analysis. Ellipses show SE.

The ratio of photosynthesis to water use is known as water use efficiency and was investigated given the changes in *A* and RWC. A life-time integrated measure of water use efficiency is carbon isotope discrimination. This was determined for grain collected from plants grown under Terminal Drought and control conditions and presented as delta (Δ). Δ is calculated using the carbon isotope ratio of both the plant samples and of air (Farquhar *et al*., 1989), and is positive in nature; typically C3 plants have a Δ of approximately 20 × 10^-3^ or 20‰. Two-way ANOVA found no significant interaction between the genotypes and the water regime, but both water regime and genotype effects were significant and large (F_(1,29)_ = 147.26, *p <* 0.001, 7^2^= 0.84 (partial) and F_(5,29)_ = 10.05, *p <* 0.001, 7^2^ = 0.63 (partial), respectively). Drought increased Δ by approximately 2.45 compared to watered control. Under both conditions, line SAL2.1 and a double mutant composed of line SAL1.1XSAL2.1 (DM) both have significantly lower Δ than the Chara control. Additionally, under drought line SAL1.2 also has a significantly reduced Δ (Table 1). In summary, given the lack of difference in g_s_ combined with a reduction in Δ levels, we conclude improved maintenance of photosynthetic capacity in TaSAL mutants under water-limited conditions.

**Table 1:**
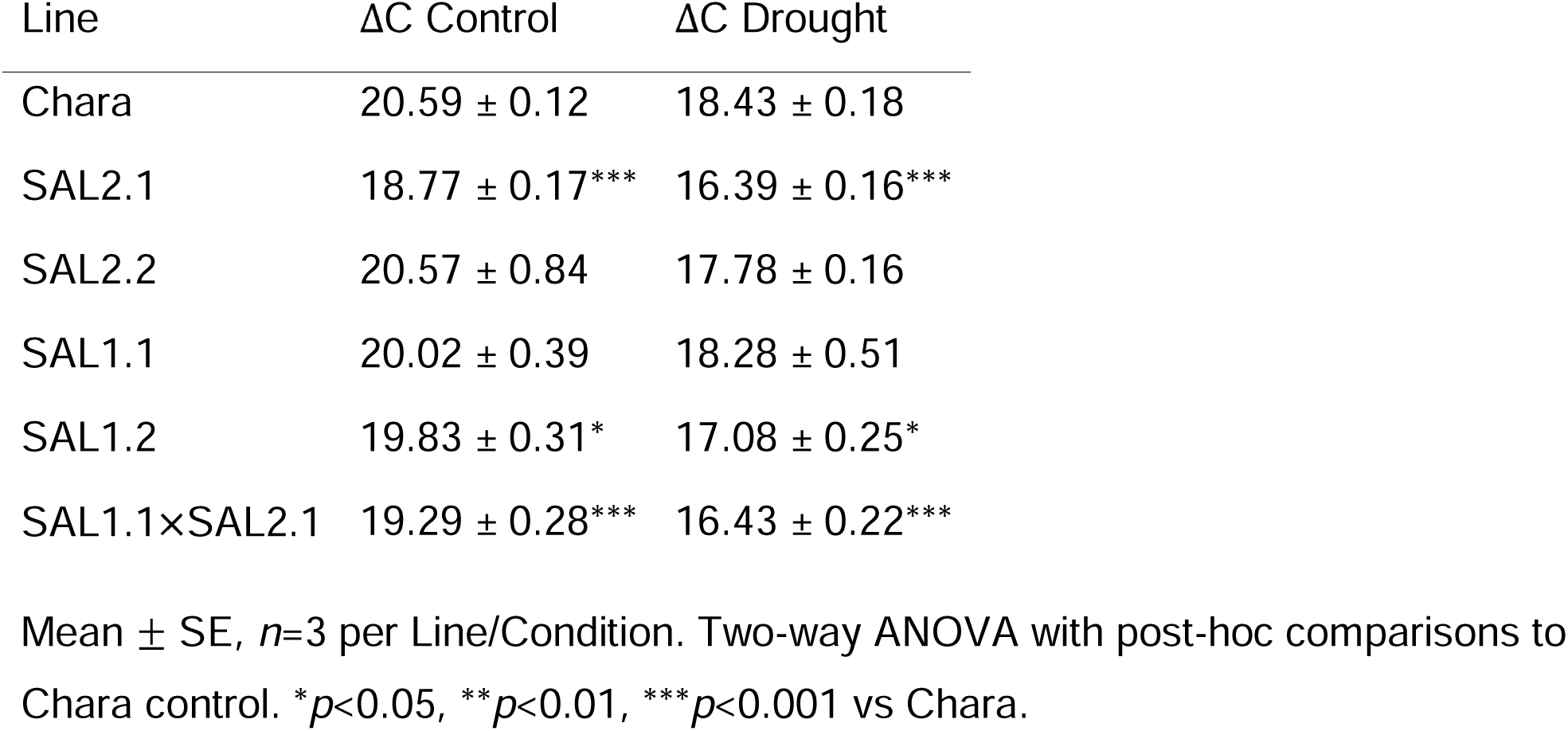
Carbon isotope discrimination (L1C) in glasshouse conditions.

We then measured ABA content of leaf material under both control and Terminal Drought conditions on all single mutants to evaluate if there is any interaction between ABA and PAP (Supporting Information Fig. S7). A three-way repeated measures ANOVA was performed to evaluate the interaction of growth conditions and ABA content, using paired sampling at days 0 and 28 of the experiment. There was a statistically significant three-way interaction between genotype, treatment and time, (F_(5, 40)_ = 3.163, *p =* 0.017, 7^2^= 0.16). At 0 days, interaction of genotype and treatment was not significant, nor treatment, but some significance was applied to genotype alone (F_(1, 40)_ = 3.71, *p <* 0.05); conversely, genotype, treatment and their interaction were all significant. Simple-simple main effects of genotype were found to be significant under drought, and pairwise comparison to Chara control, using a Holm correction, found all lines had significantly lower ABA content after 28 days drought. We identified that the dilution of TaSAL had the effect of inhibiting the ABA increase due to drought seen in the Chara control line. Critically, since all genotypes depleted soil water at equivalent rates, this reduced ABA accumulation reflects enhanced stress tolerance rather than stress avoidance, indicating that TaSAL mutants require lower stress hormone concentrations to maintain physiological function under identical water limitation conditions. Of interest is the wide variance in values for Chara, implying a higher stress susceptibility compared to the single mutant lines. This observation is consistent with an hypothesis of PAP-mediated resilience limiting stress sensitivity and consequential ABA induction in the mutant lines.

### Examination of retrograde signaling in field environments

Field trials were conducted at four distinct agroecological sites, Birchip, Yanco, Condobolin, and Narrabri, between 2015 and 2019 (Figure 4). The sites encompass essentially the breadth of the cereal growing regions of south-east Australia, separated by ∼1,000 km. The sites have varied soil types, average rainfall (decreasing with latitude from 550 to 320 mm per year) and temperature profiles (from average monthly highs of 28.9 to 24.3°C) (Summary Tables S3 to S7). The sites were used over five years, resulting in large seasonal variation. See Methods for details on field management and trial types. Collectively, this enabled assessment in 15 different growth environments spanning years of both high and low water availability, temperature and light. For example, actual total water availability ranged from 190 mm to 725 mm across the growing seasons; actual October average temperatures during grain fill ranged from 21 to 30°C for highs, and establishment lows for April ranged from 6 to 14°C (Figure 4, Supporting Information Tables S3-S7).

**Figure 4.**
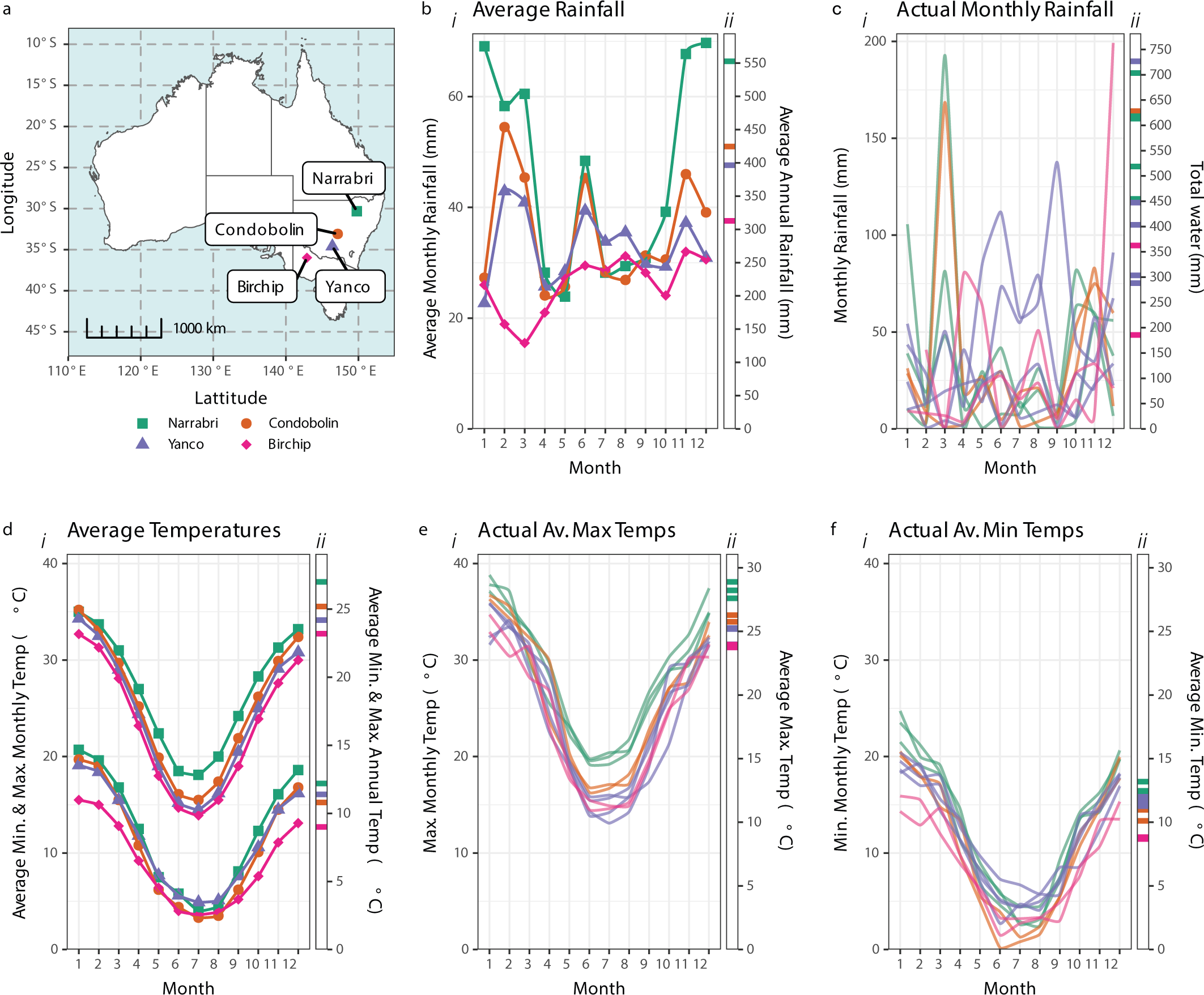
Trial environmental characteristics. (a) Trial locations across southeastern Australia. (b) Average rainfall patterns: (i) monthly averages and (ii) annual totals by site. (c) Actual rainfall variation: (i) monthly rainfall by trial and (ii) total water availability per trial. (d-f) Temperature patterns: (i) average monthly temperatures and (ii) actual temperature ranges across trials, showing (d) overall temperature patterns, (e) maximum temperatures, and (f) minimum temperatures.

Field-based phenological assessments revealed no substantive changes in growth and morphology relative to the parent line, Chara. The one exception was line SAL2.1 (the line with substantive off-target deletions) and its associated double mutants, which showed a shorter and spindly stature compared to Chara and the other two TaSAL2 deletion alleles, consistent with it having more off-target lesions (Supporting Information Dataset 1).

To assess stomatal function in field conditions, we measured canopy temperature as a proxy for stomatal aperture. Canopy temperature reflects stomatal aperture, which PAP can directly and rapidly regulate (Wilson *et al*., 2009; Estavillo *et al*., 2011; Pornsiriwong *et al*., 2017). Exogenous application of PAP closes wheat and barley stomata within minutes, as rapidly as ABA (Pornsiriwong *et al*., 2017; Zhao *et al*., 2019), though PAP acts through Respiratory Burst Oxidase Homolog D (RBOHD) rather than RBOHF for apoplastic ROS production (Tee *et al*., 2025). We measured canopy temperatures six times over seven hours of daylight. Measures were collected at Narrabri, NSW, on two consecutive days in 2017 and once in 2019 using infrared cameras deployed on drones and helicopters.

TaSAL mutant canopy temperatures showed distinct diurnal patterns compared to Chara (Figure 5, Supporting Information Fig. S8 and S9). In the morning, both 2017 and 2019 trials showed no significant differences between any genotypes and Chara, indicating no constitutive alterations in stomatal aperture under cooler conditions.

**Figure 5.**
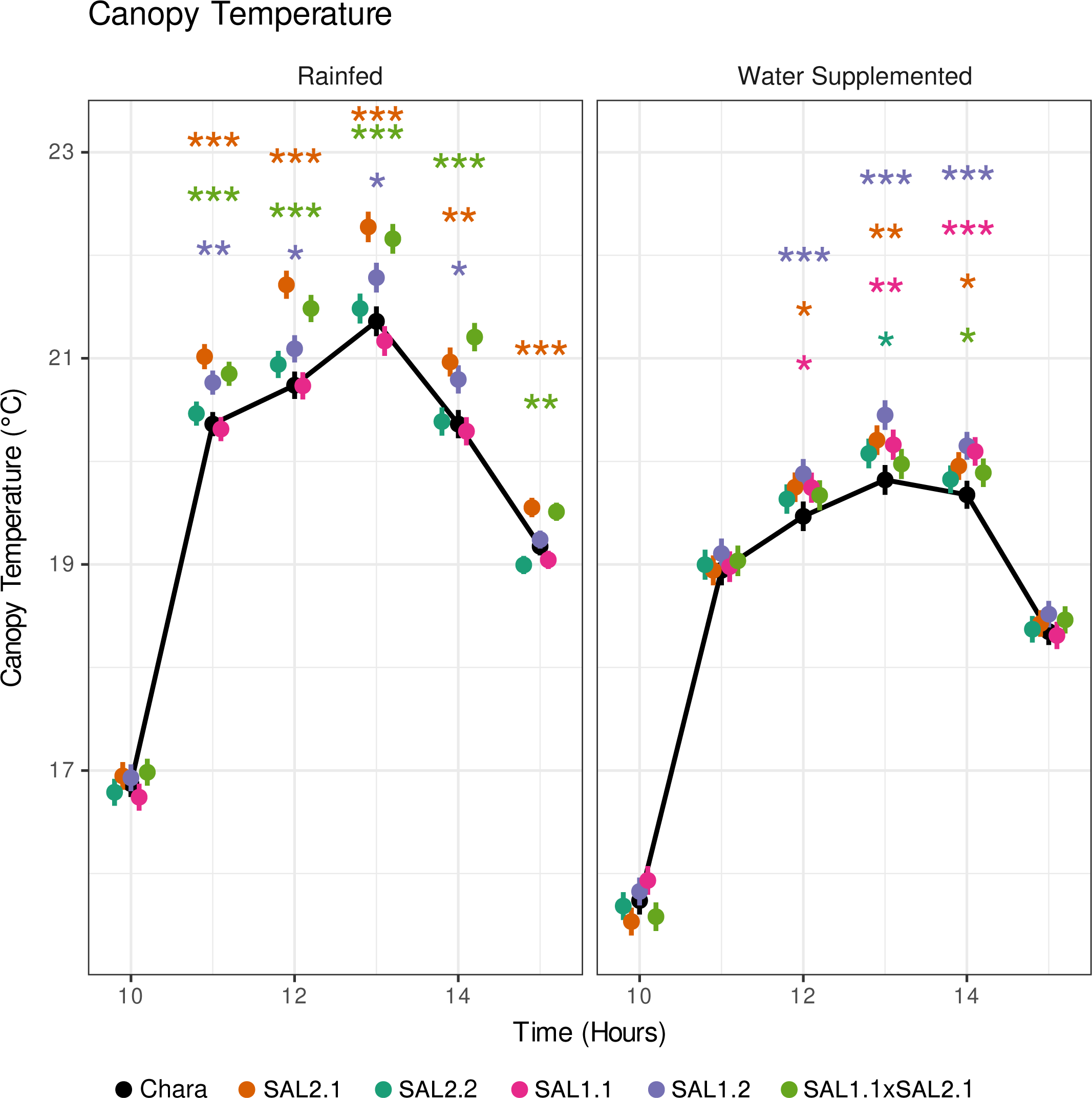
Canopy temperature, Narrabri 2017. Diurnal canopy temperature patterns measured using UAV-based infrared cameras under rainfed and water-supplemented conditions. TaSAL mutants showed increased midday temperatures vs Chara, indicating enhanced stomatal closure during peak evapotranspiration demand. Multivariate repeated measures linear mixed effects model. Values are BLUEs ± SE. \**p <* 0.05, \*\**p <* 0.01, \*\*\**p <* 0.001.

During the middle of the day, however, TaSAL lines showed significantly increased canopy temperatures compared to Chara, indicating greater stomatal closure. In 2017 trials, the pattern of significance differed between water treatments: under rainfed conditions, lines SAL2.1 and the SAL1.1XSAL2.1 double mutant showed significant increases, while under water-supplemented conditions, lines SAL1.2, SAL2.1, and SAL1.1 were highly significant, with SAL2.2 and the SAL1.1XSAL2.1 double mutant also showing significance. Temperature increases reached up to 1.2°C above Chara. In 2019 trials, conducted under higher ambient temperatures (maximum 27-31°C vs 19-22°C in 2017), only lines SAL2.1 and the SAL2.3XSAL1.1 double mutant showed significant increases at 1:15 pm under irrigated conditions, though the absolute temperature differences were greater under rainfed conditions where significance was not achieved. Significantly, the increased temperature/greater closure during the mid-day period is consistent with an enhanced environmental sensitivity - more protective of water losses - than exhibited by the controls. By the afternoon/evening, temperature differences between genotypes returned to non-significant levels in both trial years.

This pattern is consistent with the lines having enhanced environmental sensitivity rather than baseline stomatal defects, differing from prior publications in pot experiments for complete knockout/down lines. The enhanced midday stomatal closure suggests increased sensitivity to peak evapotranspiration conditions when demand is highest.

Photosynthesis-related traits were predicted for field grown materials using hyperspectral imaging, enabling the measurement of photosynthetic parameters at scale. Hyperspectral data was collected for the two trials at Narrabri 2019 (Supporting Information Fig. S10). Predictions of photosynthesis-related traits were produced using the Wheat Physiology Predictor (Furbank *et al*., 2021). This predicts leaf mass per area (LMA), nitrogen content per unit area (N_area_) and mass (N_mass_), maximum rate of carboxylation of photosynthesis at ambient temperature (V_cmax_), 25°C (V_cmax25_) and corrected for nitrogen (V_cmax25_/N_area_, electron transport rate (*J*), CO_2_ assimilation rate (Photo), chlorophyll content (SPAD) and stomatal conductance (g_s_/Cond).

Linear mixed effects models found no changes in V_cmax_ measures in either trial, with the exception of one double null (SAL1.2XSAL2.2) in rainfed conditions, and another double SAL2.3XSAL1.1 when expressed as V_cmax25_/N_area_. Maximum rates of carboxylation were more stable under water supplementation for those lines than rainfed, but no large differences could be identified for these traits. Leaf mass and nitrogen per unit area were increased in line SAL1.2, or double nulls derived from line SAL1.2 under rainfed conditions. These lines also exhibited an increased SPAD/chlorophyll content. One line, SAL1.2XSAL2.3, demonstrated a consistent increase in stomatal conductance, but no other lines exhibited this change. Overall, photosynthesis was not constitutively altered in ambient conditions, consistent with no constitutive changes in PAP levels.

### Analysis of life-time integrated measures of retrograde signaling

Yield, biomass and water productivity were assessed as life-time integrated traits of plant performance, reasoning that if the effect of repeated transient differences in retrograde signaling is cumulative, altered PAP responsiveness will be reflected in the alteration of these traits in different environments, compared to Chara (Figure 4).

These data were assessed by a two-stage multi-environment trial (MET) analysis of 15 trials that varied substantially for temperature, rainfall, irrigation or lack thereof, and latitude. We used a Factor Analytic approach (Smith *et al*., 2001; Gezan *et al*., 2022). As expected, variation between environments impacted yield, biomass and water productivity (individual trial BLUEs presented in Figure 6; Supporting Information Table S8); more interestingly, yield differences between SAL genotypes and Chara exhibited considerable variability. For example, line SAL2.2 had a range of yield differences, from an increase of over 30% in Condobolin 2017, down to a low of −7% in very hard conditions in Birchip 2018.

**Figure 6.**
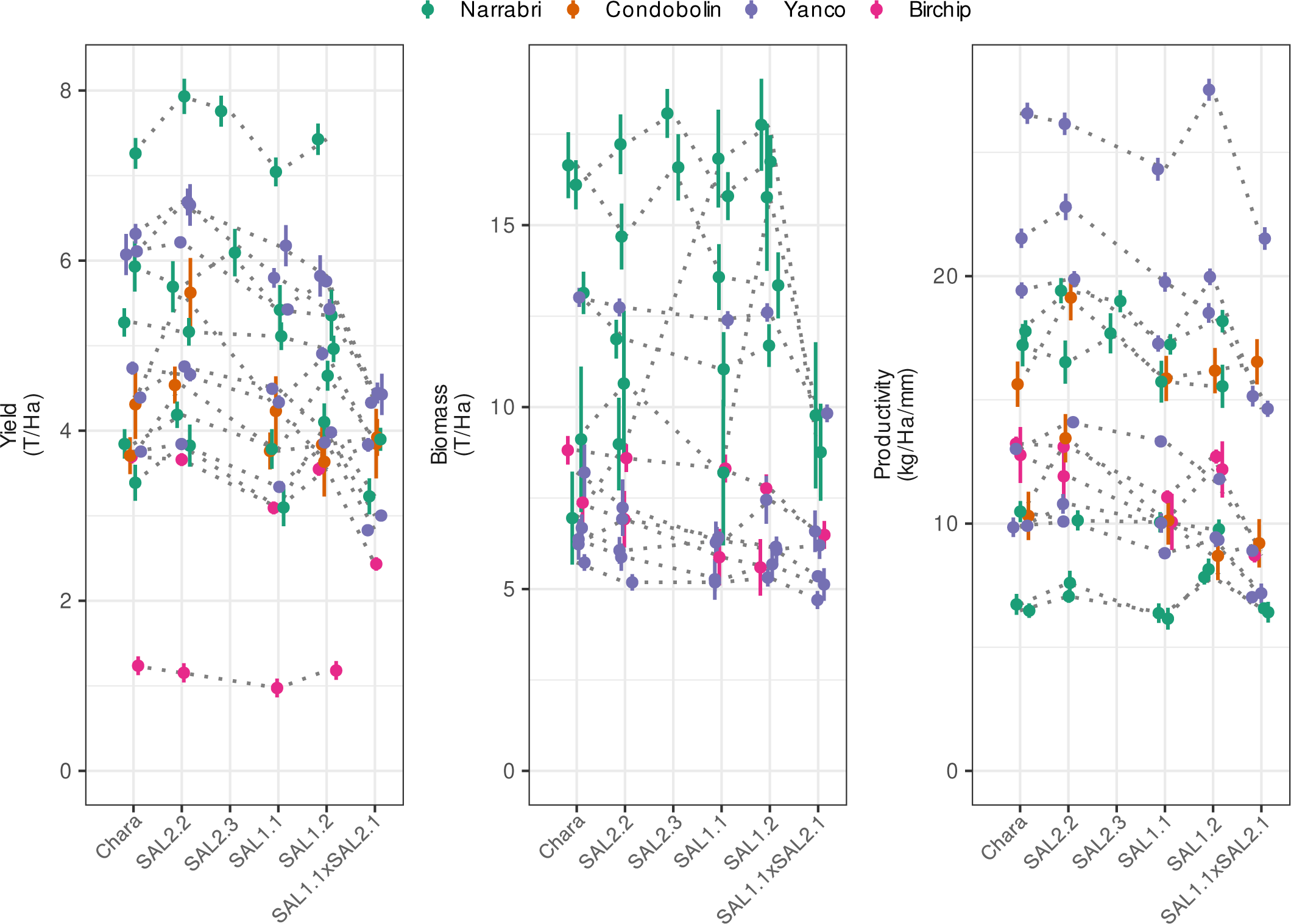
Field trial performance of TaSAL mutants. BLUEs (Best Linear Unbiased Estimators) for yield, biomass, and water productivity across trials. TaSAL2 lines outperformed Chara; TaSAL1 lines showed reduced performance.

Predictions of yield averaged across trials found one line, SAL2.2, had a significant increase in yield compared to control (approximately 4%), while its sister, SAL2.3, demonstrated an increased yield of 8%. In the integrated MET analysis this was not significant, due to the small number of trials it was included in, however in those trials it was grown, there was a statistically relevant increase when analysed as single sites (Table 2). In contrast to the TaSAL2 yield gain, the TaSAL1 subtype had a consistent yield penalty compared to Chara. (Table 2).

**Table 2:** Yield, Biomass and Water Productivity Best Linear Unbiased Predictions (BLUPs)

| Genotype | Yield (t/Ha) | Biomass (t/Ha) | Water Productivity (kg/Ha/mm Water) |
| --- | --- | --- | --- |
| Chara | 4.70 ± 0.03 | 9.41 ± 0.11 | 14.20 ± 0.10 |
| SAL2.2 | 4.89 ± 0.03*** | 9.38 ± 0.11 | 14.34 ± 0.10*** |
| SAL2.3 | 5.09 ± 0.20† | 9.83 ± 0.46 | 14.29 ± 0.51† |
| SAL1.1 | 4.46 ± 0.03*** | 9.69 ± 0.11 | 13.28 ± 0.10*** |
| SAL1.2 | 4.57 ± 0.03*** | 9.78 ± 0.11* | 13.83 ± 0.10*** |
| SAL1.1×SAL2.1 | 3.85 ± 0.03*** | 8.02 ± 0.14*** | 11.68 ± 0.10*** |

**Notes:** Mean values of random effects ± SE. \* $p < 0.05$ , \*\* $p < 0.01$ , \*\*\* $p < 0.001$ vs Chara. †Not significant in MET model, but significant in trials present.

A multivariate 3rd order Factor Analytic model was used to combine these results with appropriate weighting, allowing direct prediction of best linear unbiased predictions (BLUPs) for all lines and to predict stability of yield across sites. FA1 accounted for 90% of the variance. Together the three factors in the analytic model accounted for approximately 95% of all variance across the environments. Model biplot indicating cultivar (points) suitability to environments (arrows) and genetic covariance, indicating the magnitude of genetic variation in each environment and the degree of genetic correlation between environments, is included in Supporting Information Fig. S11. We tested the model using environmental factors, such as latitude, rainfall, irrigation, and temperature, via PCA or Random Forests and they did not explain FA1; rather, it was best explained by the genotypes, namely deletions in SAL homologues.

Water productivity measures yield per unit of actual available water for a given site’s growing season (kg/Ha/mm). Line SAL2.2 showed higher water productivity compared to the Chara control, demonstrating its improved yield given the same availability of water (Table 2). Line SAL2.3 also showed improved water productivity compared to Chara, though this improvement was not statistically significant in the MET analysis, but was for the individual sites. Again, the TaSAL1 subtype mutants showed significantly poorer water productivity compared to the Chara control and the TaSAL2 types.

Biomass was also measured in select field environments. Unlike yield and water productivity, few differences in biomass were observed. Line SAL1.2 had a significant increase compared to Chara, while the SAL1.1XSAL2.1 double mutant was smaller. Overall, no patterns could be seen in biomass changes between lines and the Chara control.

Stability of yield is a measure of the amount of variance of yield across different environments for a given genotype. It was calculated using the Superiority measure (P_i_) of Lin & Binns (1988), using the mean square difference between a cultivar’s yield and the maximum yield in each environment. This measure is less affected by unbalanced trials than other stability measures. The results indicated that the stability of the TaSAL2 null lines has improved compared to the Chara control, whereas the TaSAL1 lines have had a decrease in stability (Table 3). This could reflect improved environmental sensing for TaSAL2 and impaired sensing for TaSAL1 lines compared to Chara.

**Table 3:** Cultivar superiority (P*_i_*) estimates for yield.

| Entry | Stability ($P_i$) |
| --- | --- |
| SAL2.3 | 0.272 |
| SAL2.2 | 0.488 |
| Chara | 0.706 |
| SAL1.2 | 0.927 |
| SAL1.1 | 1.087 |
| SAL1.1×SAL2.1 | 2.362 |

Main lines investigated only included. Smaller values indicate greater stability of yield across environments.

## Discussion

### Enhanced Crop Performance Through Precision Stress Signaling

There has been a view in crop improvement that there is often an inherent trade-off between yield potential and stress resilience. Development of high-yielding varieties can introduce environmental yield penalties or instability through inadvertent selection under optimal conditions (Dietz *et al*., 2021). For example, lettuce breeding led to shallower root systems that perform well under irrigation but poorly in stress (Jackson, 1995). In wheat, such trade-offs manifest in multiple traits including nutrition, flowering time, and architecture (Dwivedi *et al*., 2021). Even traits selected specifically for stress tolerance can penalize yield - as seen with stem carbohydrate storage, which improves drought survival but reduces grain-filling efficiency in good seasons (Blum, 1998; Rebetzke *et al*., 2008).

Our findings indicate that this perceived trade-off can be overcome through precisely modulating stress signaling pathways to enhance both productivity and resilience.

Rather than constitutive changes that improve stress tolerance at the cost of growth, we created wheat lines with primed stress responses - plants that detect and respond to environmental stress earlier and more sensitively, without developmental costs. This enhanced environmental sensing translated to improved yield and water productivity, water use efficiency and sustained photosynthesis under stress conditions. This tolerance rather than avoidance mechanism provides the mechanistic basis for the enhanced field performance observed across diverse water-limited environments.

To assess if retrograde signaling variation was functioning at field scale, we used canopy temperature as a proxy for stomatal responses. TaSAL mutants showed increased canopy temperatures specifically during midday periods, with no differences morning or evening, indicating enhanced rather than constitutive regulation (Figure 5). Temperature differences were more pronounced under rainfed versus irrigated conditions (approaching 1°C), consistent with primed lines dynamically regulating stomatal aperture to optimise photosynthesis and water conservation.

Across 15 diverse field environments, wheat lines with targeted reductions in specific TaSAL2 gene copies consistently outperformed the parent cultivar. Such lines showed yield improvements of 4-8% - a substantial gain considering current yield improvement rates. Critically, these same lines also demonstrated enhanced water productivity and stress tolerance.

However, the precision of genetic targeting proved crucial. While lines with TaSAL2 deletions consistently improved, those with TaSAL1 deletions showed yield penalties, demonstrating that enhanced performance requires precise pathway modulation rather than broad disruption of stress signaling. The contrasting effects of TaSAL1 versus TaSAL2 deletions highlight that targeted modification of specific loci, rather than complete pathway disruption, is essential for achieving these benefits.

### Priming Environmental Responsiveness

Knockouts of TaSAL2 or TaSAL1 did not lower SAL activity or increase PAP in unstressed conditions, but in response to water deficit did lower relative SAL activity in most lines by up to 60% and could elevate PAP levels. The mild attenuation of SAL activity is expected to allow PAP levels to have a greater capacity to rise early in stress. In turn, this will lead to a modified pattern of stress signals then communicated to the nucleus. What is significant is that titrating SAL activity raises PAP only under stress, thereby generating a set of lines that prime retrograde signal responsiveness.

Compared to *Arabidpsis sal1* knockout mutants and previous wheat studies that knocked out all or most TaSAL genes (Rossel et al., 2006; Wilson et al., 2009; Pornsiriwong et al., 2017; Mohr et al., 2022; Abdallah et al., 2022), our primed, selective knockdown lines exhibited minimal growth phenotypes during glasshouse evaluations (Supporting Information Fig. S4). Under both watered and water-deficit conditions, growth, tiller numbers, and plant height remained largely consistent with the Chara control, suggesting that partial TaSAL reduction avoids the developmental penalties associated with complete pathway disruption whilst enabling enhanced stress responses. The primed lines demonstrated improved water use efficiency, maintained relative water content, and sustained photosynthetic rates under drought conditions.

Most significantly, deletion of either TaSAL subtype enabled continued photosynthetic assimilation during water limitation - demonstrating the benefits of selective over complete pathway disruption (Figure 3, Supporting Information Fig. S6). This maintenance of photosynthesis reveals the sophisticated interplay between chloroplast-mediated sensing and ABA-independent PAP regulation of stomatal function (Pornsiriwong et al., 2017), allowing dynamic rather than constitutive stress responses.

The molecular basis for this enhanced environmental sensing lies in PAP’s dual role as both a stress signal and stomatal regulator (Pornsiriwong *et al*., 2017; Tee *et al*., 2025). PAP accumulates rapidly under oxidative stress when SAL enzyme activity is reduced (Estavillo et al. 2011, Chan et al. 2016a). In our partially knocked-down lines, this established mechanism creates a more sensitive environmental detection system. Unlike constitutive stress responses, this priming mechanism is expected to allow PAP levels to rise earlier and reach higher peaks during stress episodes, enabling plants to detect and respond to environmental perturbations before damage occurs.

The transient nature of PAP accumulation in primed lines reflects the sophisticated regulation of this pathway. Previous work in *Arabidopsis* demonstrated that approximately 80% reduction in AtSAL1 activity is required before significant PAP accumulation occurs (Phua et al., 2018a); our knockdown lines operate far below this level. This creates a dynamic system where early PAP increases likely trigger rapid stress responses (Estavillo et al., 2011) that could subsequently restore SAL activity through feedback mechanisms, potentially suppressing further PAP accumulation. Such regulatory feedback loops explain why PAP increases appear modest in single-timepoint measurements despite the enhanced stress responsiveness observed in our physiological assessments.

The equivalent water depletion rates across all genotypes during controlled drought experiments definitively establish that enhanced performance results from improved stress tolerance rather than reduced water consumption. This tolerance mechanism explains the paradoxical observation of reduced ABA accumulation in TaSAL mutants under drought: rather than indicating reduced stress exposure (avoidance), the lower ABA levels reflect enhanced cellular resilience that reduces the need for ABA-mediated stress responses. PAP acts directly on stomatal guard cells, bypassing traditional ABA signaling pathways (Pornsiriwong et al., 2017), enabling more efficient stress responses that maintain cellular function without requiring elevated stress hormone concentrations.

This tolerance rather than avoidance mechanism identified under controlled conditions provides the mechanistic basis for the enhanced field performance observed across diverse water-limited environments. PAP-mediated stomatal control enables dynamic water management throughout the day. Field measurements of canopy temperature revealed that primed TaSAL lines increased stomatal closure specifically during midday stress periods while maintaining normal aperture during benign morning and evening conditions. This temporal regulation may optimise the trade-off between CO_2_ uptake for photosynthesis and water conservation, explaining the observed improvements in both water use efficiency and sustained photosynthetic rates under water limitation.

### Gene-Specific Engineering Enables Targeted Pathway Modulation

The two homeologous groups were found to be the result of a conserved duplication that we conclude has occurred twice in the Poaceae. The conservation of the duplication in the Pooideae, distinct tissue expression and co-expression analysis patterns, along with contrasting effects in field assessments of knockouts of TaSAL1 and TaSAL2 subtypes, are evidence towards specialization within subgroups; however, the nature of this subgrouping is far from clear, be it simply distinct expression levels or reflects subfunctionalisation, which may be suggested by the different co-expression profiles for TaSAL1 and TaSAL2.

Co-expression analysis provided molecular evidence for functional specialisation between TaSAL homeologue groups. TaSAL1 homeologues showed highest expression in vegetative tissues and were enriched for fundamental cellular processes including aminoacyl-tRNA biosynthesis and porphyrin metabolism, consistent with their high expression during active vegetative growth phases (Soprano *et al*., 2018; Adjei *et al*., 2023). In contrast, TaSAL2-associated networks showed stronger enrichment for metabolic flexibility pathways including amino acid metabolism, energy metabolism, and secondary metabolism (phenylpropanoid biosynthesis).

Potentially, this molecular evidence could support the divergence of function between gene families, with TaSAL1 specialized for supporting vegetative growth and TaSAL2 for environmental stress adaptation. These pathway enrichments align with established roles of branched-chain amino acid degradation (Pires *et al*., 2016), amino acid metabolism coordination (Joshi *et al*., 2010; Bowne *et al*., 2012; Yadav *et al*., 2019), and phenylpropanoid biosynthesis (Dong & Lin, 2021) in plant adaptation to variable water availability.

### From Transient Signals to Lifetime Performance

Traditionally, operational retrograde signaling has been studied under controlled conditions, revealing regulation of oxidative stress genes over seconds to hours (Chan *et al*., 2010; Crawford *et al*., 2018). However, chloroplasts continuously sense environmental conditions, and constitutive alterations to retrograde signals lead to sustained developmental changes, as in the case of, B-cyclocitral (Jiang & Dehesh, 2021; Sierra *et al*., 2022). This raises a fundamental question: can transient retrograde signals that operate on minutes-to-hours timescales accumulate to influence plant performance across an entire growing season? If so, primed retrograde responsiveness should manifest as altered performance differences across diverse field environments, indicating integration of multiple environmental fluctuations, rather than a forgotten response to single acute events.

Critically, the fact that both homeologous groups show altered stability relative to Chara - TaSAL2 beneficial and TaSAL1 detrimental - demonstrates that priming the responsiveness of retrograde signaling has successfully altered environmental responses across the entire growing season (Table 3). This interpretation is further supported by our Factor Analytic model, where FA1 accounted for 90% of the variance across environments, strongly suggesting that the specific TaSAL gene modifications are fundamental to explaining the differential environmental responses observed. This differential environmental responsiveness across multiple field seasons represents the strongest evidence that these lines are truly primed for enhanced retrograde signaling, with the direction of the stability change determined by the specific functional role of each homeologous group. The contrasting stability profiles demonstrate that targeted modification of specific TaSAL genes has altered how plants integrate and respond to environmental information across their lifecycle, validating our hypothesis that retrograde signaling effects accumulate beyond transient stress responses.

## Conclusions

Modulating retrograde signaling in field environments results in locus-specific differences in biomass, yield, water productivity, and yield stability. The TaSAL1 locus improves resilience at the expense of yield, whereas the TaSAL2 locus results in better resilience, 4-8% gains in yield and thus improved yield stability. Therefore, improving the dynamic nature of retrograde signaling, such as the priming of the SAL-PAP pathway herein, can enhance responsiveness to dynamic environments by maintaining photosynthesis whilst acclimating to transient stress and water deficits. Critically, we show that retrograde signals integrate environmental information throughout the plant lifecycle, not just during acute stress episodes. This precision approach to pathway modulation demonstrates that selective gene targeting can develop climate-adapted crops without yield penalties.

## Acknowledgements

The authors acknowledge and thank Dr Zhongyi Li, formerly of CSIRO Agriculture & Food, for access to the HIB population and for his advice. We also thank Dr Hilary Stuart-Williams of the Farquhar Laboratory and ANU’s Stable Isotope Laboratory for assistance and guidance of o^13^C isotope discrimination analysis. We thank Professor Owen Atkin for his helpful discussion and encouragement. This work was supported by the Australian Grains Research and Development Corporation (GRDC) through grant ANU00020. AFB, MM and BJP were supported by an Australian Research Council Australian Laureate Fellowship (FL190100056). BJP, KXC and ZP are supported by the Australian Research Council Training Centre for Accelerated Future Crops Development (IC210100047).

## Author Contributions

AY, GJR, GME and BJP conceived the experiment(s), MM, AY, JZ, MDM, ZP, EET, EKA, CAH and DPC conducted the experiment(s), AFB, MM and BJP analysed the results. AFB, MM, KXC and BJP wrote and reviewed the manuscript. BJP and AY share corresponding authorship. AFB, MM and AY contributed equally to this work.

## Data Availability

The data that support the findings of this study are available in the Supporting Information of this article. Genotyping data are provided in Supporting Information Dataset 1, field trial environmental characteristics are detailed in Supporting Information Tables S3-S7, and comprehensive yield, biomass and water productivity data across all trials are presented in Supporting Information Table S8. Additional raw phenotypic measurements are available from the corresponding authors upon request for research purposes.

## Supporting information

Additional supporting information may be found in the online version of the article at the publisher’s website.

**Dataset 1** Genotype-by-sequencing results for all mutants in study.

**Fig. S1** Comparison of variation in wheat SAL gene expression across developing seedling leaves.

**Fig. S2** Representative agarose gel of screening for TaSAL mutants via PCR.

**Fig. S3** PAP content under irrigated and drought conditions in glasshouse experiments.

**Fig. S4** Plant height and relative water content under drought.

**Fig. S5** Soil weight changes during drought

**Fig. S6** Photosynthesis and stomatal conductance measures.

**Fig. S7** ABA content of glasshouse-grown wheat material under Control and Drought conditions.

**Fig. S8** Leaf/canopy temperature results from Narrabri 2019.

**Fig. S9** Leaf/canopy temperature results from Narrabri 2017.

**Fig. S10** Hyperspectral-derived estimates of photosynthesis related traits.

**Fig. S11** Factor analytic model visualization of multi-environment trial analysis - Biplot and Genetic Covariance Matrix heatmap.

**Table S1** Table of primers used in discovery of *TaSal* null lines from HIB population.

**Table S2** Line identifiers and *TaSal* gene knockout.

**Table S3** Narrabri environmental characteristics for all years of trials.

**Table S4** Condobolin environmental characteristics for all years of trials.

**Table S5** Yanco environmental characteristics for all years of trials.

**Table S6** Birchip environmental characteristics for all years of trials.

**Table S7** Characterising average and actual water conditions per site and year, including irrigation where used.

**Table S8** Yield, Biomass and Water Productivity Best Linear Unbiased Predictions for all trials.

**Method S1** Phylogenetic analysis.

**Method S2** PAP extraction, preparation, and quantification by HPLC.

