## Supporting Information for "Modulation of SAL retrograde signaling promotes yield and water productivity responses in dynamic field environments"

### New Phytologist Supporting Information



**Article acceptance date:** 2 September 2025


**The following Supporting Information is available for this article:**


**Fig. S1** Comparison of variation in wheat SAL gene expression across developing seedling leaves.

**Fig. S2** Representative agarose gel of screening for TaSAL mutants via PCR.

**Fig. S3** PAP content under irrigated and drought conditions in glasshouse experiments.

**Fig. S4** Plant height and relative water content under drought.

**Fig. S5** Soil weight changes during drought.

**Fig. S6** Photosynthesis and stomatal conductance measures.

**Fig. S7** ABA content of glasshouse-grown wheat material under control and drought conditions.

**Fig. S8** Leaf/canopy temperature results from Narrabri 2019.

**Fig. S9** Leaf/canopy temperature results from Narrabri 2017.

**Fig. S10** Hyperspectral-derived estimates of photosynthesis-related traits.

**Fig. S11** Factor analytic model visualisation of multi-environment trial analysis.

**Table S1** Primers used in discovery of *TaSAL* null lines from HIB population.

**Table S2** Line identifiers and *TaSAL* gene knockouts.

**Table S3** Narrabri environmental characteristics for all trial years.

**Table S4** Condobolin environmental characteristics for all trial years.

**Table S5** Yanco environmental characteristics for all trial years.

**Table S6** Birchip environmental characteristics for all trial years.

**Table S7** Average and actual water conditions per site and year, including irrigation.

**Table S8** Yield, biomass and water productivity best linear unbiased predictions for all trials.

**Methods S1** Phylogenetic analysis.

**Methods S2** PAP extraction, preparation and quantification by HPLC.

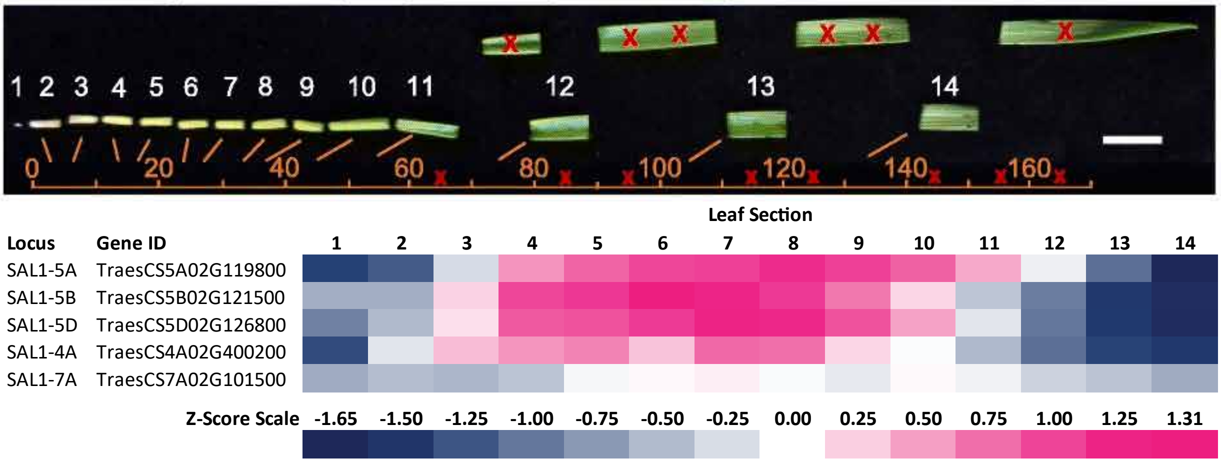


Fig. S1: **Individual SAL expression levels vary by gene and location in leaf material.** SAL gene IDs (TraesCS5A02G119800, TraesCS5B02G121500, TraesCS5D02G126800, TraesCS4A02G400200, and TraesCS7A02G101500) were searched in RNA-seq data generated by Loudya et al., 2021. Z-scores indicate variation in each transcript across all section of a 6-day old seedling. Figure was adapted from Loudya et al. (2021) (scale bar 100 mm).

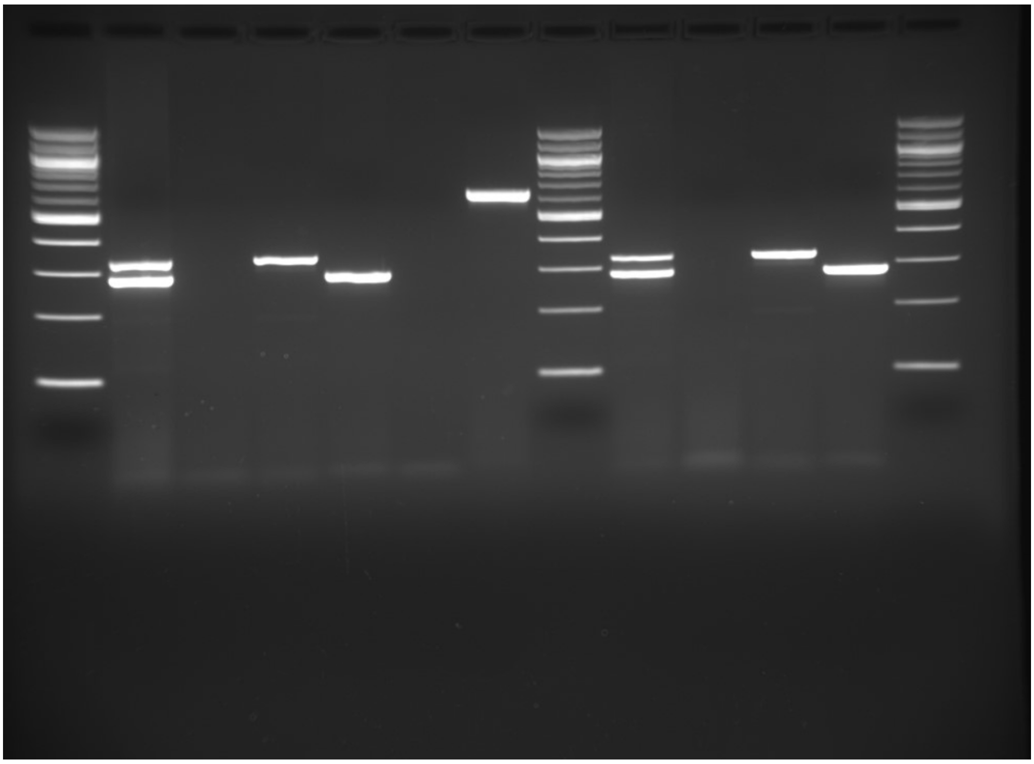

Fig. S2: Representative agarose gel of screening for TaSAL1 mutants. *Lanes 1,8,13* 100bp Ladder (Biolab), *2* Chara, *3* DM (SAL1.1×SAL2.1), *4* SAL2.1 (4A), *5* SAL1.1 (5D), *6* No Template Control, *7* DM (SAL1.1×SAL2.1) with TaSAL1 7A Primers (Positive PCR Control), *9* Chara, *10* DM (SAL1.1×SAL2.1), *11* SAL2.1 (4A), *12* SAL1.1 (5D). All lanes except *7* used both TaSal1 5D and 4A primer sets.

A

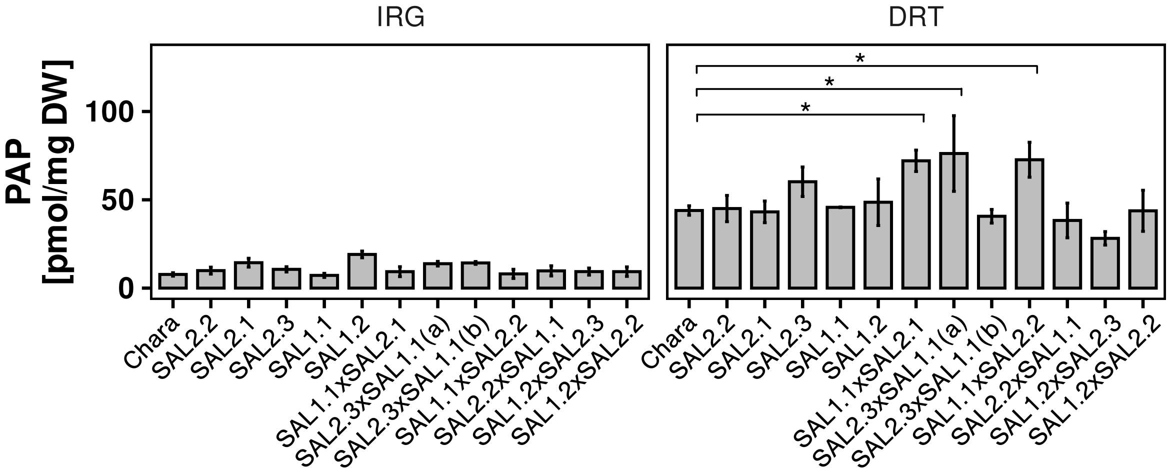


Fig. S3: **PAP content (A) under Irrigated and Drought conditions in glasshouse experiments.** Only significant differences can be seen under drought conditions, where PAP content is generally higher than control, suggesting a priming of the SAL1-PAP retrograde signalling pathway, increasing its responsiveness, is possible in these lines. Genotype $\times$ treatment interaction: PAP content $F_{(12,52)} = 2.42, p = 0.014$. The main effect for PAP content under stress was genotype ($p < 0.001$ $F_{(12,52)} = 4.61$. False discovery rate (FDR) correction was applied to comparisons with the Chara control.

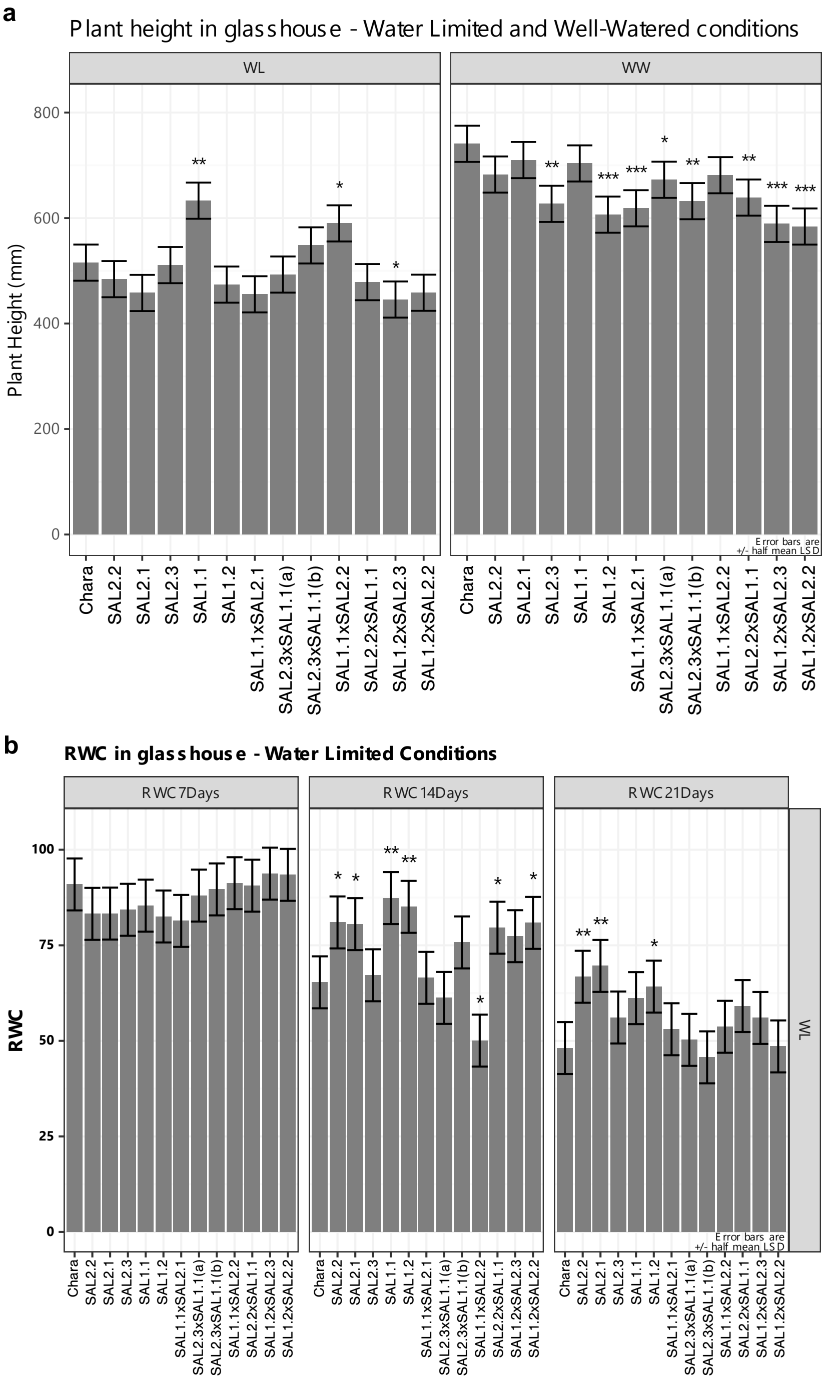


Fig. S4: Plant height and relative water content under drought. (A) Plant height of Chara and *TaSal1* lines under control (WW) and water limited (WL) conditions in glasshouse experiment. Linear mixed models were used to account for spatial variation in glasshouse. Wald test indicated that Genotype and Condition interaction was significant ($F_{(12,35.4)} = 0.5$, $p$=0.0178), as were Genotype ($F_{(12,35.9)} = 7.1$, $p<0.001$) and Condition ($F(1,5.5) = 361$, $p<0.001$) alone. Water limited conditions reduced stature by 149 mm ($\pm$8.3 SE). Under water limited conditions, few differences are seen, and only one double mutant is shorter than the control Chara variety. In contrast, multiple lines are shorter than control under well watered, control conditions. Values are BLUEs, significant differences indicated are to Chara control only, error bars are half least significant differences. (B) Relative eater content. Time series was modelled using a multivariate model for repeated measures in ASREML-R; No significant differences in relative water content could be seen at Day 7 of drought, but most single nulls can be seen to maintain water content at Day 14 and (less so) at Day 21. In general, singel nulls have greater RWC in drought than control or double null lines. *$p<0.05$, **$p<0.01$, ***$p<0.001$.

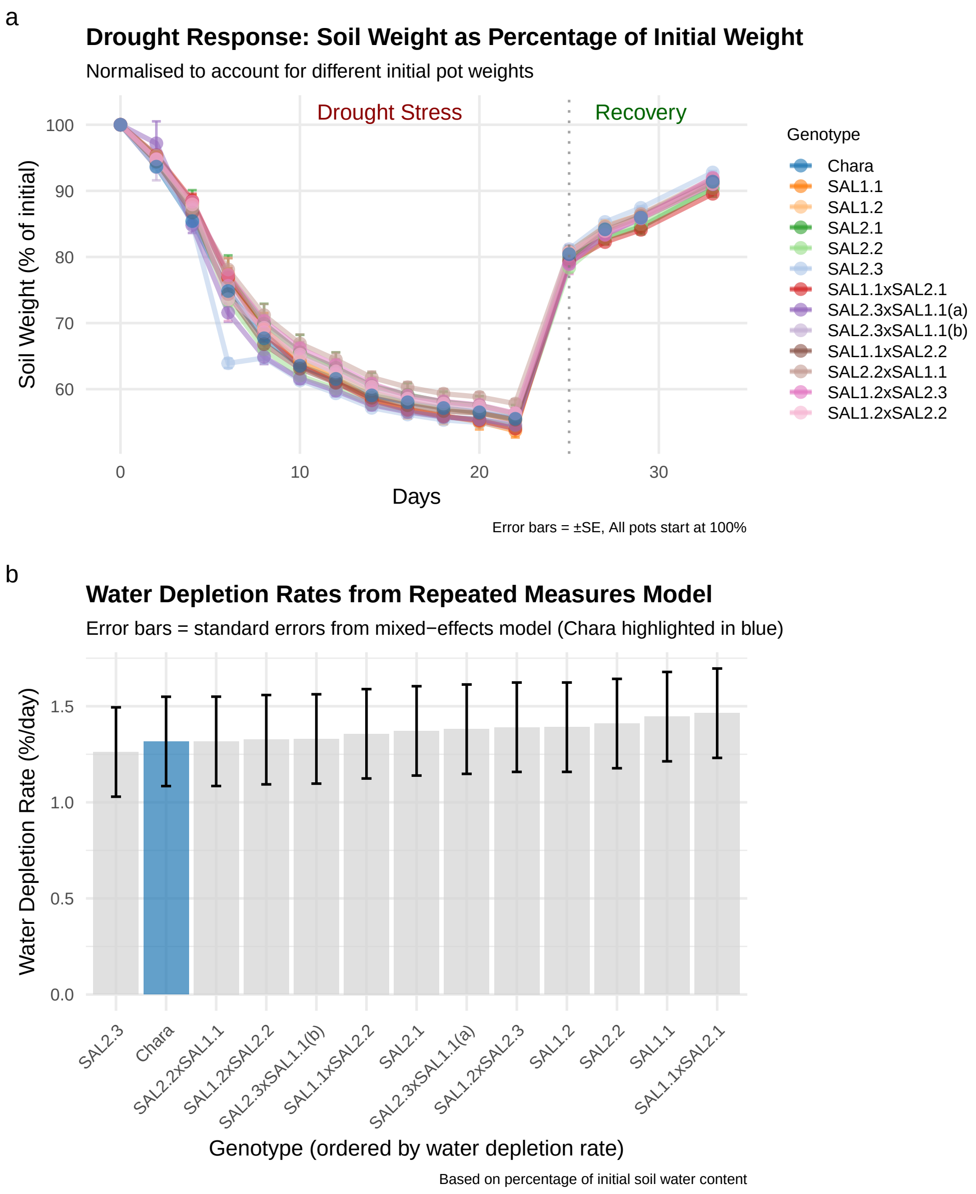


Fig. S5: **Soil weight changes during drought**. a) No significant differences in soil (expressed as percentage of initial pot weight) were found using a repeated measures mixed effects model for any lines during the onset and recovery of drought. b) daily water depletion rates of all lines (calculated as percentage mean reduction in soil weight during drought onset); significant differences between genotypes was identified. Chara control indicated in blue.

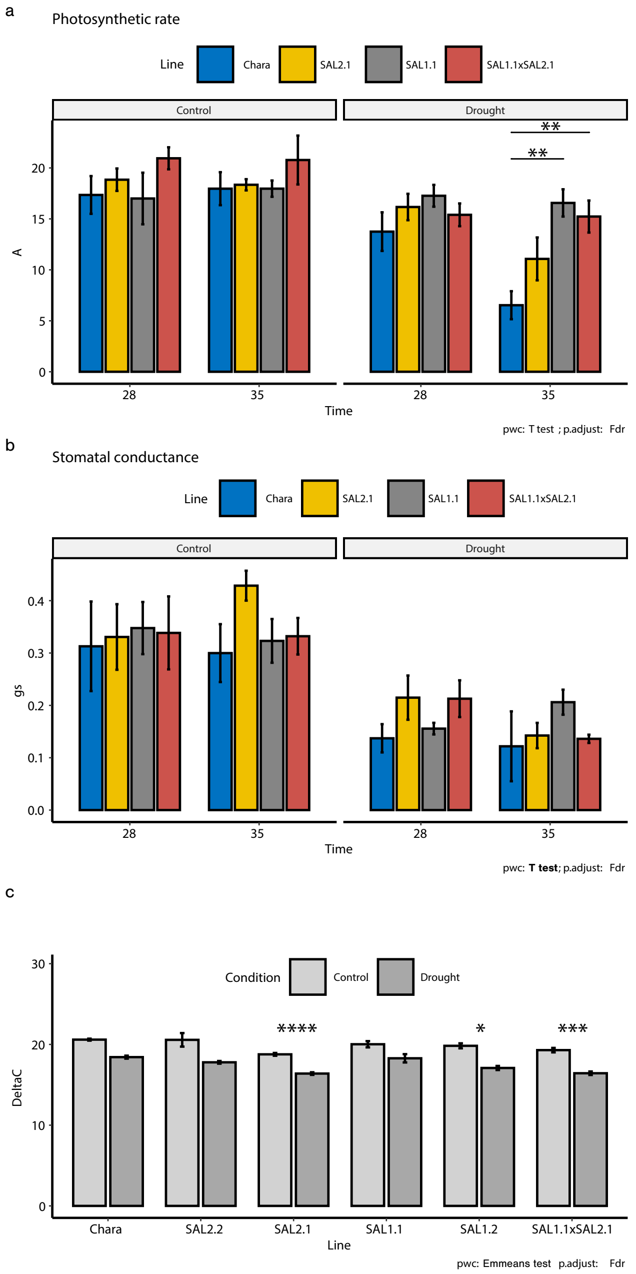


Fig. S6: **Photosynthesis, stomatal conductance and carbon isotope discrimination (Δ).**. A) Assimilation rate: Mixed effects model showing significant genotype effect (F(3,11) = 6.67, p = 0.008) with TaSAL deletion lines maintaining higher rates under drought compared to Chara. B) Stomatal conductance: Mixed effects model showing drought reduced conductance across all lines (F(1,24) = 89.25, p < 0.0001, $\eta^2_G$ = 0.53) with no genotype-specific differences. Error bars show SE. Drought conditions significantly reduce stomatal conductance in all lines, but no significant differences between Chara and mutants were identified. C) ΔC13: No significant differences in interaction between genotypes and drought were found, but main effects for both were apparent. Drought significantly increased ΔC13 compared to control conditions, and genotypes SAL2.1, SAL1.2 and the double mutant (SAL1.1xSAL2.1) were all significantly higher in ΔC13 than Chara control, indicating an increase in WUE in these lines in all conditions. ***p*<0.05, ***p*<0.01, ****p*<0.001, *****p*<0.0001.

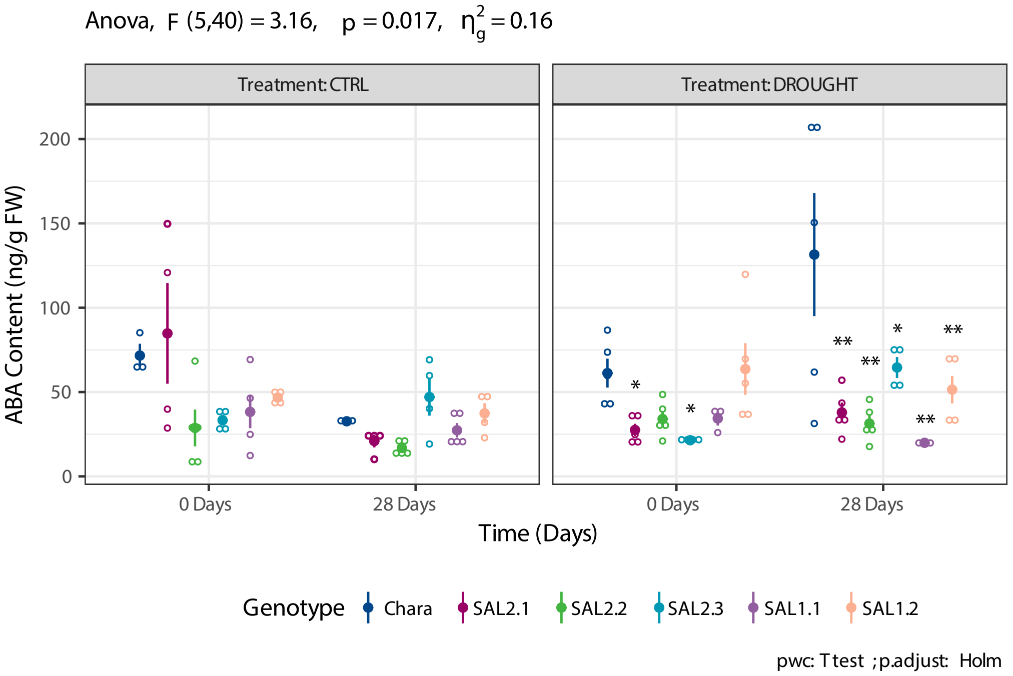


Fig. S7: ABA content of glasshouse grown wheat material under Control and Drought conditions, on day 0 and day 28 of treatment. Three-way repeated measures ANOVA. Individual points and Mean ± SE indicated. There is a statistically significant three-way interactions between Treatment, Genotype and Time, $F_{(5, 40)} = 3.163$, p = 0.017. There was a statistically significant simple simple main effect of Genotype on ABA content for Drought conditions at both timepoints ($p<0.05$ for all), but not for Control treatment. Pairwsie T-Tests were completed for all compared to Chara, using a Holm correction. Note, several extreme outliers were removed from the Control treatment set by flagging data points that fall below Q1 - 3IQR or above Q3 + 3IQR, where Q1 and Q3 are the first and third quartiles, and IQR is the interquartile range. These samples' removal has reduced the effectiveness of the control treatment batch to identify any Day 0 differences. * $p < 0.05$, ** $p < 0.01$; n=5 per genotype per treatment.

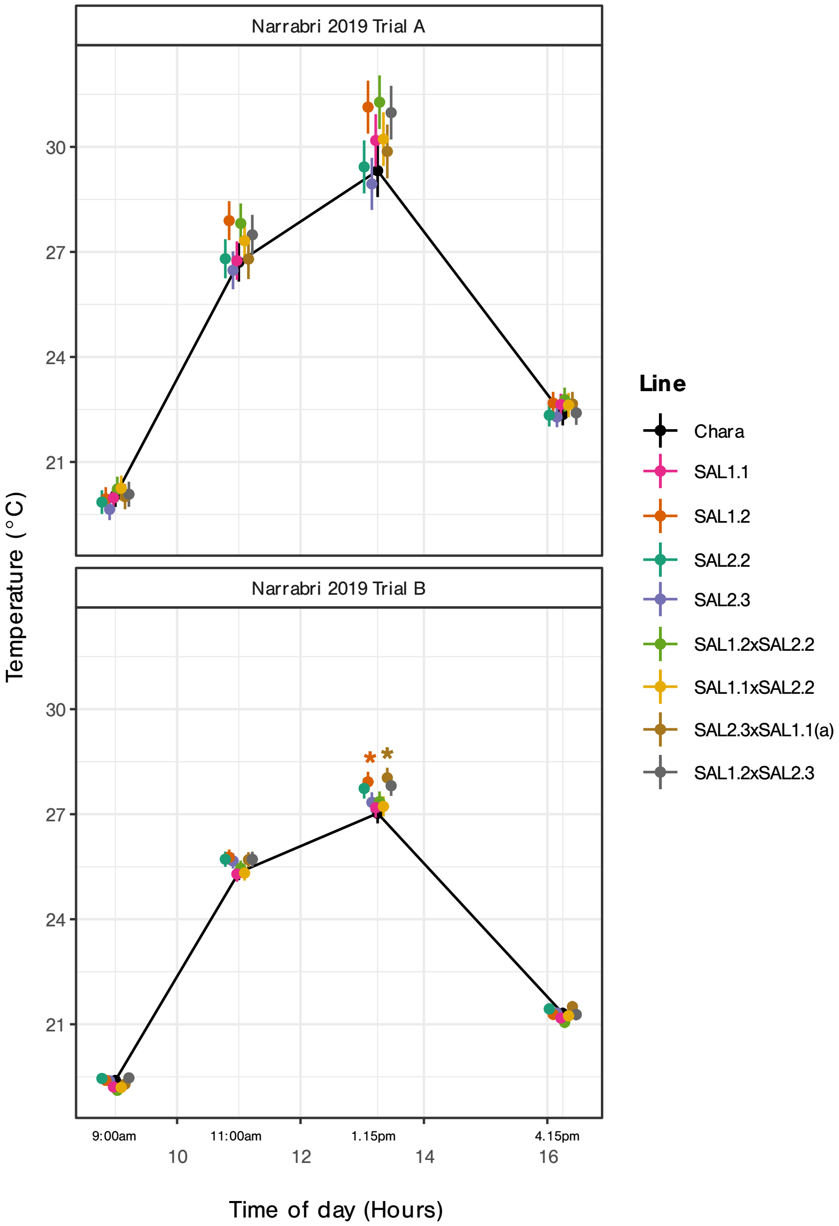


Fig. S8: **Leaf temperature results from two field trials measured at four time points during one day.** Narrabri 2019 Trial A was grown under a non-watering regime, while Trial B was supplemented with some irrigation. Multivariate linear mixed models were applied to each trial result to model as a repeated measures analysis. Two lines had significantly higher leaf temperatures compared to Chara control during the 1.15pm time measurement in Trial B (SAL1.2 and SAL2.3xSAL1.1), but no other differences were identified. See Methods section for details of trial management; error bars are Standard Errors.

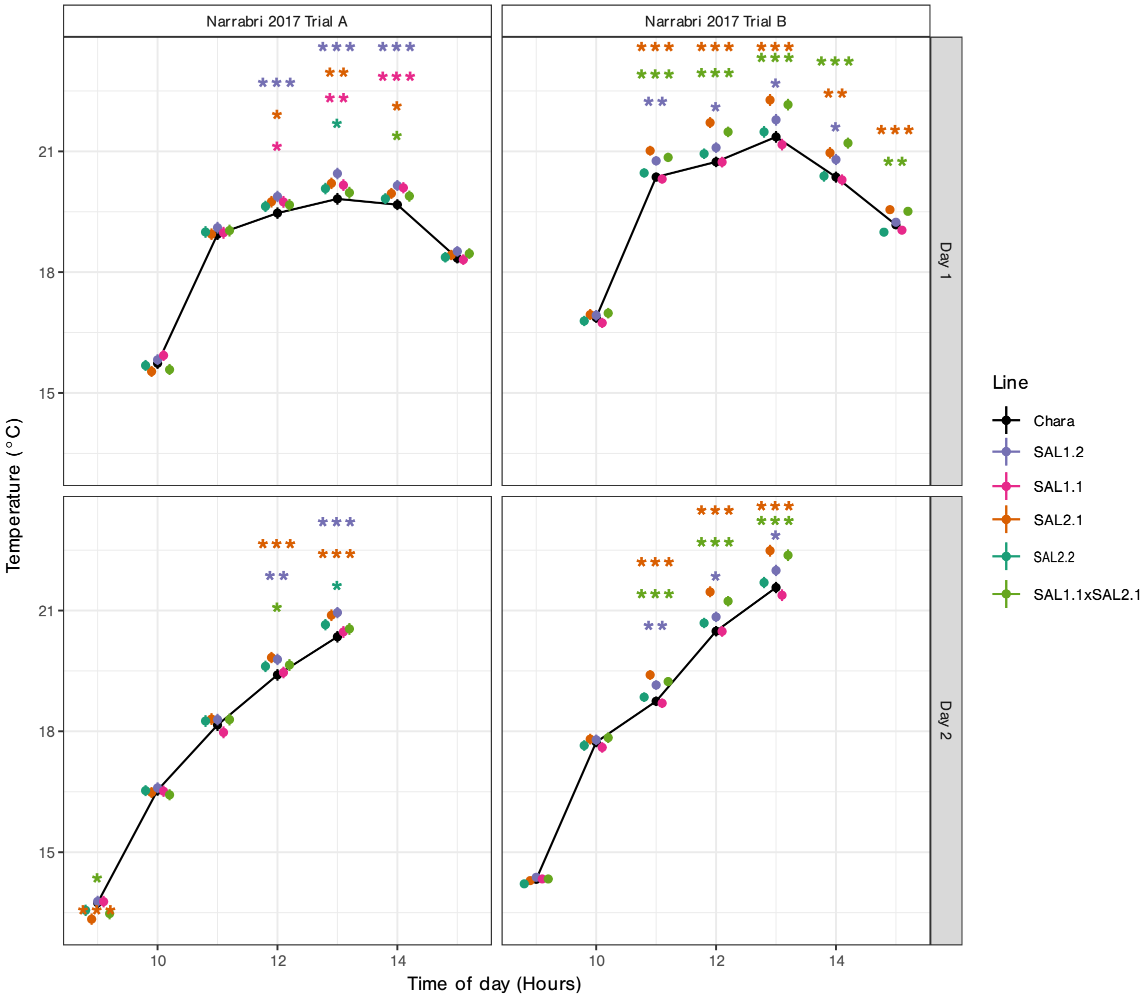


Fig. S9: **Leaf temperature results from two field trials measured between 9am and 3pm on successive days.** Narrabri 2017 Trial A was grown under a water-supplemented regime, while Trial B was not supplemented with any additional watering. Multivariate linear mixed models were applied to each trial result to model as a repeated measures analysis. Significantly increased leaf temperatures are seen in both trials and both days, but with a greater leaf temperature difference in the rainfed Trial B, which saw three lines with consistently higher temperatures in lines SAL2.1, SAL1.1xSAL2.1 and SAL1.2 (up to 0.9°C higher than Chara control). See Methods section for details of trial management; error bars are Standard Errors.

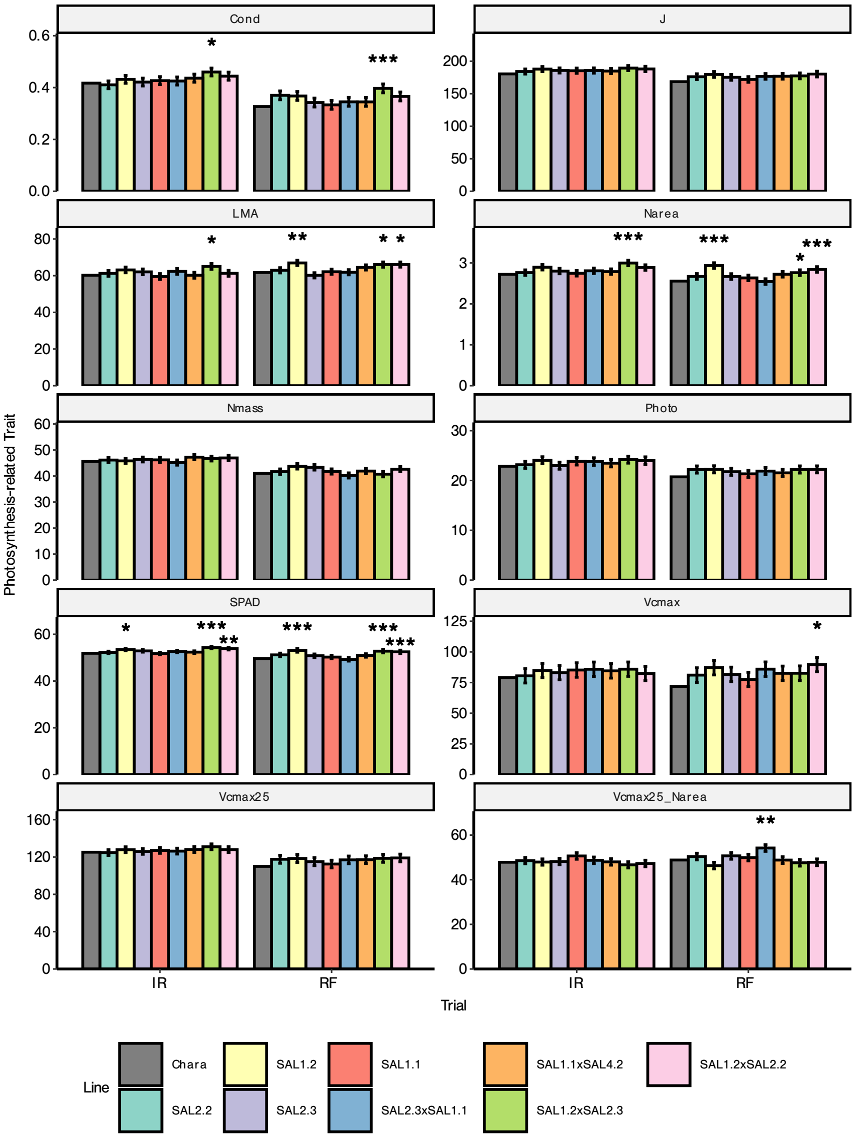


Fig. S10: Hyperspectral-derived estimates of photosynthesis related traits. Estimates of stomatal conductance (Cond, mol $H_2O$ m$^{-2}$ s$^{-1}$), election transport rate ($J$), leaf mass per unit area (LMA, g m$^{-2}$), nitrogen content per unit area ($N_{area}$) and mass ($N_{mass}$), $CO_2$ assimilation rate (Photo, $\mu$mol $CO_2$ m$^{-2}$ s$^{-1}$), chlorophyll content (SPAD), and maximum rate of carboxylation of photosynthesis at ambient temperature ($V_{cmax}$, $\mu$mol $CO_2$ m$^{-2}$ s$^{-1}$), 25°C ($V_{cmax25}$, $\mu$mol $CO_2$ m$^{-2}$ s$^{-1}$) and corrected for nitrogen ($V_{cmax25}/N_{area}$, $\mu$mol $CO_2$ s$^{-1}$(g N$^{-1}$)). Measurements are in one trial under normal rainfall conditions (RF) and under a water supplementation (IR). Error bars are standard errors calculated from marginal contrast analysis of differences between lines and Chara control from linear mixed effects models. Stars represent significant differences compared to Chara control line. *$p < 0.05$, **$p < 0.001$, ***$p < 0.0001$.

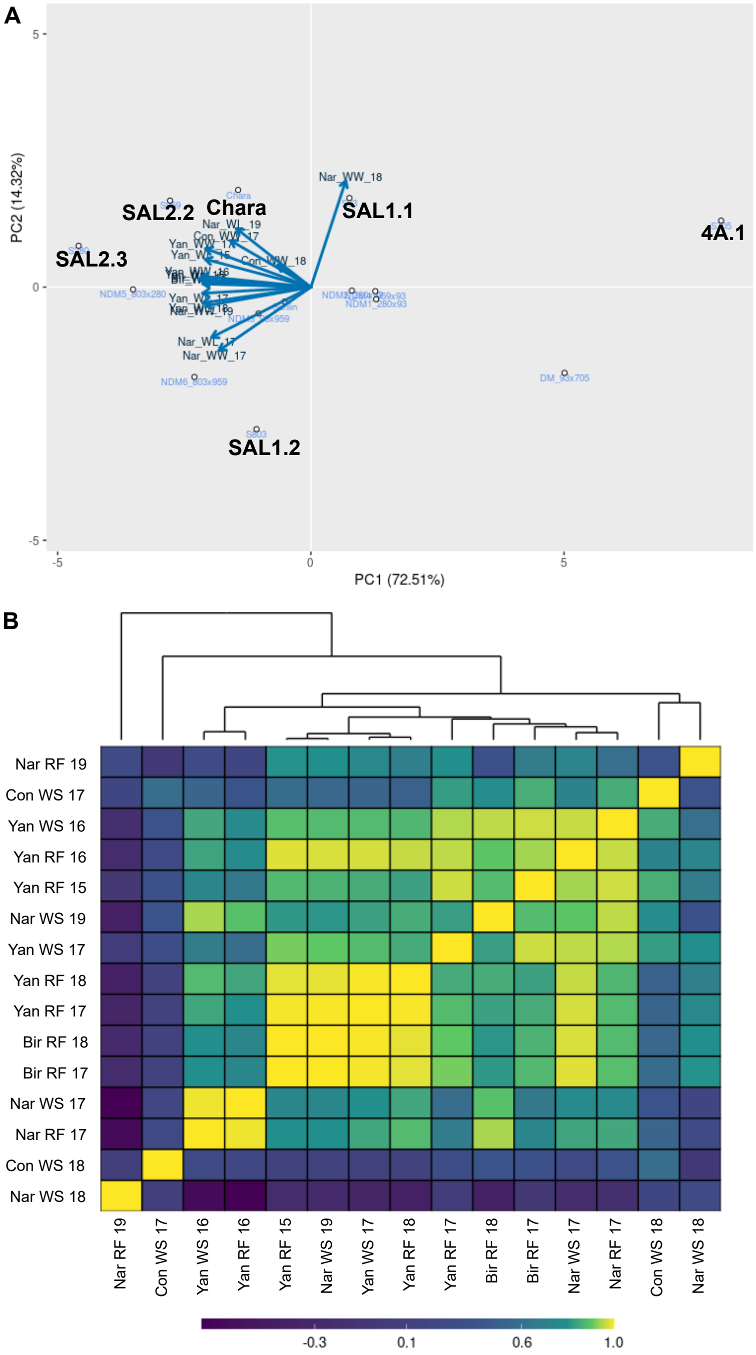


Fig. S11: Factor analytic (FA) model visualization of multi-environment trial data. (A) Biplot displaying the first two factors from a factor analytic model fitted to yield data across environments. Environment vectors (arrows) indicate the strength and direction of environmental responses, where vector length represents the amount of genetic variance explained in each environment, and angles between vectors indicate genetic correlations between environments (acute angles suggest positive correlations, obtuse angles suggest negative correlations). Genotype scores (points) show the relative performance and stability of varieties, where points closer to the origin indicate stable performance and points further from the origin show specific environmental adaptation. (B) Genetic covariance matrix heatmap derived from the FA model, showing the pattern and magnitude of genetic variances (diagonal) and covariances (off-diagonal) between environments. Lighter colours indicate stronger positive genetic covariances, suggesting similar genetic responses between environment pairs. This matrix decomposition provides insights into the complexity of genotype-by-environment interactions and helps identify environment groups with similar genetic responses.

Table S1: Table of primers used in discovery of *TaSal1* null lines from HIB population.

| Target gene | 5' primer sequence | 3' primer sequence |
|---|---|---|
| *TaSal1* 5D | CCCACCATTACTCTACCATCTA | ATGCTTCACTAAACCAGTTATGC |
| *TaSal1* 4A | GGCACCACACTACTGCCTTC | GTGTTCACGGTTAATCAGTACA |
| *TaSal1* 7A | GAGTTCTGACCTATGACTCTTCA | CGGATAGTTTACATAGAGCTGGT |

Table S2: Line identifiers and *TaSal1* gene knockout.

| Line Identifier | Genotype |
|---|---|
| SAL1.1 | 5D - TaSAL1 |
| SAL1.2 | 5D - TaSAL1 |
| SAL2.1 | 4A - TaSAL2 |
| SAL2.2 | 4A - TaSAL2 |
| SAL2.3 | 4A - TaSAL2 |
| SAL1.1×SAL2.1 | Double |
| SAL2.3×SAL1.1 | Double |
| SAL2.3×SAL1.1 | Double |
| SAL1.1×SAL2.2 | Double |
| SAL2.2×SAL1.1 | Double |
| SAL1.2×SAL2.3 | Double |
| SAL1.2×SAL2.2 | Double |

Table S3: Narrabri environmental characteristics

| Narrabri | Rainfall (mm) | | | | | | | | | | | | |
|---|---|---|---|---|---|---|---|---|---|---|---|---|---|
| | Jan | Feb | Mar | Apr | May | Jun | Jul | Aug | Sep | Oct | Nov | Dec | Annual |
| **2001-2020** | **69.1** | **58.3** | **60.5** | **28.2** | **23.9** | **48.4** | **28.2** | **29.4** | **30.9** | **39.2** | **67.7** | **69.7** | **552.7** |
| *Max* | *196.0* | *176.6* | *192.8* | *107.0* | *57.4* | *186.4* | *74.6* | *102.8* | *123.4* | *88.2* | *200.0* | *247.4* | *891.4* |
| *Min* | *5.0* | *3.2* | *2.2* | *0.0* | *0.0* | *0.6* | *0.6* | *1.0* | *0.2* | *3.6* | *7.0* | *6.8* | *206.2* |
| 2017 | 105.6 | 12.0 | 192.8 | 13.2 | 24.8 | 41.8 | 11.4 | 19.8 | 3.6 | 62.8 | 59.8 | 37.8 | 585.4 |
| 2018 | 38.8 | 19.0 | 48.2 | 18.0 | 0.6 | 7.6 | 7.8 | 31.2 | 9.4 | 81.6 | 58.2 | 56.0 | 376.4 |
| 2019 | 10.2 | 3.2 | 81.4 | 0.0 | 29.6 | 0.6 | 13.8 | 1.0 | 0.6 | 4.6 | 54.4 | 6.8 | 206.2 |
| | **Max Temp °C** | | | | | | | | | | | | |
| | Jan | Feb | Mar | Apr | May | Jun | Jul | Aug | Sep | Oct | Nov | Dec | Mean |
| **2001-2020** | **35.0** | **33.7** | **31.0** | **27.0** | **22.4** | **18.5** | **18.1** | **20.0** | **24.2** | **28.3** | **31.3** | **33.2** | **27.0** |
| *Max* | *38.8* | *37.1* | *33.3* | *30.0* | *23.7* | *19.8* | *20.4* | *23.1* | *27.7* | *31.2* | *35.4* | *37.4* | *28.9* |
| *Min* | *30.3* | *29.9* | *29.0* | *24.0* | *21.0* | *15.3* | *16.4* | *17.2* | *20.3* | *24.7* | *27.4* | *28.0* | *25.2* |
| 2017 | 37.8 | 37.1 | 29.7 | 25.5 | 23.1 | 19.2 | 19.2 | 20.5 | 25.9 | 28.9 | 29.5 | 34.8 | 27.6 |
| 2018 | 37.1 | 34.9 | 33.0 | 29.7 | 23.3 | 19.8 | 20.4 | 20.7 | 25.1 | 28.8 | 31.0 | 34.9 | 28.2 |
| 2019 | 38.8 | 35.7 | 32.9 | 28.4 | 22.3 | 19.6 | 20.0 | 21.8 | 26.7 | 30.3 | 32.7 | 37.4 | 28.9 |
| | **Min Temp °C** | | | | | | | | | | | | |
| | Jan | Feb | Mar | Apr | May | Jun | Jul | Aug | Sep | Oct | Nov | Dec | Mean |
| **2001-2020** | **20.7** | **19.6** | **16.8** | **12.5** | **7.5** | **5.8** | **4.0** | **4.5** | **8.1** | **12.3** | **16.1** | **18.6** | **12.2** |
| *Max* | *24.7* | *21.2* | *19.1* | *14.6* | *9.9* | *7.7* | *6.1* | *6.3* | *9.6* | *14.6* | *19.6* | *20.6* | *13.2* |
| *Min* | *17.3* | *17.4* | *13.8* | *9.6* | *3.3* | *3.9* | *1.3* | *2.3* | *6.5* | *9.9* | *13.3* | *16.0* | *11.2* |
| 2017 | 23.5 | 21.2 | 18.3 | 11.3 | 8.3 | 6.0 | 2.6 | 3.2 | 7.2 | 13.6 | 14.3 | 19.7 | 12.4 |
| 2018 | 21.5 | 19.1 | 18.1 | 14.6 | 6.8 | 4.6 | 2.9 | 2.5 | 9.3 | 14.2 | 16.3 | 19.7 | 12.5 |
| 2019 | 24.7 | 20.0 | 19.1 | 13.6 | 9.0 | 5.8 | 4.5 | 4.5 | 7.5 | 13.5 | 15.5 | 20.6 | 13.2 |

Table S4: Condobolin environmental characteristics

| **Condobolin** | **Rainfall (mm)** | | | | | | | | | | | | |
|---|---|---|---|---|---|---|---|---|---|---|---|---|---|
| | **Jan** | **Feb** | **Mar** | **Apr** | **May** | **Jun** | **Jul** | **Aug** | **Sep** | **Oct** | **Nov** | **Dec** | **Annual** |
| **2000-2020** | **27.3** | **54.5** | **45.4** | **24.1** | **25.7** | **45.3** | **28.3** | **26.9** | **31.3** | **30.6** | **46.0** | **39.1** | **424.6** |
| *Max* | *90.0* | *172.1* | *168.5* | *136.8* | *95.4* | *161.5* | *59.7* | *70.7* | *143.2* | *76.9* | *108.6* | *111.8* | *698.6* |
| *Min* | *1.3* | *0.0* | *0.2* | *0.2* | *0.4* | *4.4* | *1.5* | *3.4* | *0.8* | *0.0* | *2.8* | *2.8* | *155.3* |
| 2017 | 31.2 | 4.6 | 168.5 | 20.8 | 27.0 | 4.8 | 19.0 | 20.9 | 6.2 | 52.7 | 75.0 | 59.9 | 490.6 |
| 2018 | 28.6 | 8.4 | 1.0 | 2.4 | 15.5 | 29.7 | 1.5 | 3.8 | 8.5 | 29.9 | 83.0 | 11.8 | 224.1 |
| | **Max Temp °C** | | | | | | | | | | | | |
| | **Jan** | **Feb** | **Mar** | **Apr** | **May** | **Jun** | **Jul** | **Aug** | **Sep** | **Oct** | **Nov** | **Dec** | **Mean** |
| **2000-2020** | **35.2** | **33.1** | **29.7** | **25.2** | **19.9** | **16.1** | **15.5** | **17.4** | **21.9** | **26.2** | **29.9** | **32.4** | **25.2** |
| *Max* | *39.8* | *35.9* | *32.3* | *29.9* | *21.7* | *17.6* | *17.1* | *20.1* | *23.7* | *30.8* | *34.6* | *35.7* | *26.4* |
| *Min* | *30.2* | *29.1* | *27.3* | *22.1* | *17.8* | *13.7* | *13.8* | *14.7* | *18.0* | *22.3* | *27.4* | *28.5* | *23.9* |
| 2017 | 36.7 | 35.5 | 30.3 | 23.7 | 19.6 | 16.8 | 17.1 | 17.2 | 22.8 | 27.0 | 29.8 | 32.5 | 25.8 |
| 2018 | 36.3 | 34.3 | 32.3 | 29.9 | 20.4 | 16.3 | 16.7 | 18.2 | 22.1 | 27.1 | 27.8 | 33.9 | 26.3 |
| | **Min Temp °C** | | | | | | | | | | | | |
| | **Jan** | **Feb** | **Mar** | **Apr** | **May** | **Jun** | **Jul** | **Aug** | **Sep** | **Oct** | **Nov** | **Dec** | **Mean** |
| **2000-2020** | **19.7** | **19.0** | **15.5** | **10.8** | **6.2** | **4.4** | **3.3** | **3.5** | **6.2** | **10.1** | **14.5** | **16.8** | **10.8** |
| *Max* | *25.4* | *20.9* | *18.0* | *13.5* | *9.6* | *6.4* | *6.3* | *5.4* | *8.1* | *14.0* | *18.7* | *19.9* | *12.2* |
| *Min* | *15.9* | *16.2* | *13.7* | *7.1* | *3.1* | *0.2* | *0.8* | *1.4* | *3.7* | *7.7* | *11.7* | *14.0* | 9.2 |
| 2017 | 20.5 | 17.9 | 17.1 | 9.8 | 4.9 | 0.2 | 0.8 | 1.6 | 5.5 | 10.7 | 14.6 | 17.7 | 10.1 |
| 2018 | 20.1 | 18.1 | 14.8 | 13.3 | 5.6 | 3.9 | 1.3 | 2.4 | 5.5 | 12.0 | 14.9 | 19.9 | 11.0 |

Table S5: Yanco environmental characteristics

| Yanco | Rainfall (mm) | | | | | | | | | | | | |
|---|---|---|---|---|---|---|---|---|---|---|---|---|---|
| | **Jan** | **Feb** | **Mar** | **Apr** | **May** | **Jun** | **Jul** | **Aug** | **Sep** | **Oct** | **Nov** | **Dec** | **Annual** |
| **2000-2020** | **22.7** | **42.9** | **40.9** | **25.7** | **28.5** | **39.4** | **33.8** | **35.5** | **29.8** | **29.3** | **37.2** | **30.9** | **396.5** |
| *Max* | *70.6* | *170.0* | *243.0* | *113.4* | *85.2* | *111.8* | *74.0* | *79.2* | *137.6* | *127.6* | *117.2* | *91.0* | *737.6* |
| *Min* | *0.2* | *1.0* | *0.0* | *2.0* | *1.8* | *2.2* | *5.8* | *5.0* | *1.2* | *0.0* | *3.2* | *1.6* | *189.8* |
| 2015 | 43.4 | 28.2 | 0.0 | 40.8 | 14.2 | 73.2 | 54.8 | 79.2 | 22.6 | 5.6 | 22.8 | 33.6 | 418.4 |
| 2016 | 54.2 | 13.8 | 50.4 | 11.8 | 85.2 | 111.8 | 58.2 | 66.0 | 137.6 | 46.2 | 35.0 | 67.4 | 737.6 |
| 2017 | 10.0 | 13.0 | 19.4 | 23.2 | 25.0 | 2.2 | 24.0 | 33.2 | 1.2 | 28.4 | 21.4 | 91.0 | 292.0 |
| 2018 | 24.2 | 1.0 | 4.2 | 2.0 | 23.2 | 29.2 | 5.8 | 8.8 | 12.4 | 7.4 | 57.6 | 22.4 | 198.2 |
| | **Max Temp °C** | | | | | | | | | | | | |
| | **Jan** | **Feb** | **Mar** | **Apr** | **May** | **Jun** | **Jul** | **Aug** | **Sep** | **Oct** | **Nov** | **Dec** | **Mean** |
| **2000-2020** | **34.3** | **32.5** | **29.0** | **24.4** | **19.0** | **15.2** | **14.5** | **16.2** | **20.5** | **25.0** | **29.1** | **30.8** | **24.2** |
| *Max* | *39.1* | *35.1* | *31.6* | *28.8* | *20.7* | *17.3* | *16.0* | *18.7* | *23.1* | *29.1* | *33.2* | *33.0* | *25.3* |
| *Min* | *29.7* | *28.3* | *25.2* | *21.4* | *16.7* | *13.8* | *13.1* | *13.6* | *17.2* | *20.7* | *26.3* | *22.3* | *23.1* |
| 2015 | 31.6 | 34.1 | 28.9 | 22.5 | 18.8 | 14.4 | 13.1 | 14.3 | 19.0 | 29.1 | 29.7 | 32.3 | 24.0 |
| 2016 | 32.4 | 33.4 | 31.0 | 26.9 | 19.0 | 13.9 | 14.2 | 15.8 | 17.4 | 21.3 | 28.1 | 31.9 | 23.8 |
| 2017 | 35.9 | 33.4 | 31.6 | 24.4 | 19.5 | 15.9 | 16.0 | 15.9 | 21.7 | 26.0 | 30.2 | 31.7 | 25.2 |
| 2018 | 35.8 | 33.8 | 30.9 | 28.8 | 19.4 | 15.5 | 15.7 | 16.8 | 20.9 | 26.5 | 27.5 | 31.5 | 25.3 |
| | **Min Temp °C** | | | | | | | | | | | | |
| | **Jan** | **Feb** | **Mar** | **Apr** | **May** | **Jun** | **Jul** | **Aug** | **Sep** | **Oct** | **Nov** | **Dec** | **Mean** |
| **2000-2020** | **19.1** | **18.4** | **15.5** | **11.8** | **7.7** | **5.6** | **4.9** | **5.1** | **7.6** | **10.6** | **14.5** | **16.2** | **11.4** |
| *Max* | *23.5* | *20.3* | *18.0* | *14.4* | *10.4* | *8.1* | *6.7* | *6.4* | *9.7* | *13.8* | *18.3* | *18.2* | *12.1* |
| *Min* | *15.6* | *15.3* | *13.6* | *8.7* | *5.4* | *2.7* | *3.8* | *3.3* | *5.8* | *7.8* | *12.4* | *10.5* | *10.7* |
| 2015 | 18.3 | 19.2 | 14.4 | 11.4 | 8.2 | 5.0 | 4.4 | 5.7 | 5.8 | 13.8 | 14.4 | 17.8 | 11.5 |
| 2016 | 19.5 | 17.9 | 18.0 | 12.7 | 9.7 | 7.3 | 6.7 | 5.8 | 8.5 | 8.6 | 12.4 | 16.9 | 12.0 |
| 2017 | 18.6 | 17.0 | 17.0 | 11.3 | 7.0 | 2.7 | 4.7 | 4.1 | 7.4 | 11.6 | 15.6 | 18.2 | 11.3 |
| 2018 | 20.4 | 18.9 | 15.6 | 13.5 | 7.6 | 5.2 | 4.4 | 5.0 | 7.0 | 12.2 | 14.7 | 18.1 | 11.9 |

Table S6: Birchip environmental characteristics

| Birchip | Rainfall (mm) | | | | | | | | | | | | |
|---|---|---|---|---|---|---|---|---|---|---|---|---|---|
| | Jan | Feb | Mar | Apr | May | Jun | Jul | Aug | Sep | Oct | Nov | Dec | Annual |
| **2000-2020** | **26.0** | **18.9** | **15.5** | **21.0** | **27.2** | **29.5** | **28.7** | **31.2** | **28.2** | **24.1** | **32.0** | **30.6** | **312.9** |
| *Max* | *180.6* | *54.2* | *55.6* | *95.2* | *63.6* | *58.2* | *48.8* | *72.2* | *118.6* | *71.2* | *72.6* | *199.2* | *515.8* |
| *Min* | *0.0* | *0.0* | *0.0* | *0.0* | *2.4* | *1.2* | *3.2* | *4.2* | *0.0* | *0.0* | *0.0* | *0.0* | *183.4* |
| 2017 | 0.0 | 40.8 | 0.0 | 80.0 | 63.6 | 1.2 | 17.0 | 50.8 | 5.2 | 28.6 | 33.6 | 21.2 | 342.0 |
| 2018 | 9.2 | 0.0 | 6.8 | 3.8 | 22.0 | 27.4 | 15.4 | 23.6 | 0.0 | 15.0 | 7.8 | 199.2 | 330.2 |
| | **Max Temp °C** | | | | | | | | | | | | |
| | Jan | Feb | Mar | Apr | May | Jun | Jul | Aug | Sep | Oct | Nov | Dec | Mean |
| **2004-2020** | **32.7** | **31.3** | **28.1** | **23.2** | **18.0** | **14.7** | **13.9** | **15.5** | **19.0** | **23.9** | **27.6** | **30.0** | **23.2** |
| *Max* | *36.3* | *33.8* | *31.3* | *26.7* | *19.7* | *15.9* | *14.9* | *17.3* | *21.1* | *29.3* | *32.0* | *32.8* | *24.3* |
| *Min* | *30.4* | *28.0* | *25.1* | *20.1* | *16.3* | *13.2* | *12.4* | *13.6* | *16.5* | *20.4* | *25.0* | *26.6* | *22.2* |
| 2017 | 32.9 | 30.4 | 31.3 | 22.9 | 17.7 | 15.5 | 14.9 | 14.9 | 19.2 | 24.7 | 30.1 | 30.3 | 23.7 |
| 2018 | 34.7 | 31.9 | 28.2 | 26.7 | 18.4 | 14.5 | 14.6 | 15.3 | 19.7 | 24.9 | 27.0 | 31.6 | 24.0 |
| | **Min Temp °C** | | | | | | | | | | | | |
| | Jan | Feb | Mar | Apr | May | Jun | Jul | Aug | Sep | Oct | Nov | Dec | Mean |
| **2004-2020** | **15.5** | **15.0** | **12.8** | **9.2** | **6.4** | **4.0** | **3.6** | **3.9** | **5.2** | **7.6** | **11.1** | **13.1** | **9.0** |
| *Max* | *17.0* | *17.0* | *15.5* | *11.1* | *9.6* | *6.5* | *5.7* | *5.6* | *7.3* | *10.6* | *14.4* | *15.6* | *9.7* |
| *Min* | *13.8* | *12.0* | *10.5* | *7.2* | *4.1* | *1.5* | *1.8* | *2.7* | *3.0* | *6.4* | *8.5* | *11.7* | *8.0* |
| 2017 | 14.3 | 12.9 | 14.6 | 9.8 | 5.9 | 1.5 | 2.7 | 3.3 | 4.9 | 7.6 | 13.3 | 13.5 | 8.7 |
| 2018 | 15.9 | 15.5 | 12.0 | 8.9 | 6.3 | 3.3 | 3.2 | 3.3 | 3.0 | 8.6 | 10.7 | 15.3 | 8.8 |

Table S7: Characterising average and actual water conditions per site and year, including irrigation where used. Averages provided on Monthly and Total basis, including by Growth Season (GS). PS, Pre-Sowing. Type A are 'non-irrigated' unless necessary for completion of trial, and Type B are irrigated trials.

| | | | | | Monthly | | | Total | | | | | |
|---|---|---|---|---|---|---|---|---|---|---|---|---|---|
| Type | Regimes | Location | Year | Region | Av. Rain | GS Av. Rain | GS Act. Rain | GS Av. Rain | GS Act. Rain | Irrig. | Total | PS Water | PS+GS |
| A | RF | Yanco | 2015 | CW | 33.0 | 32.4 | 39.2 | 259.1 | 313.2 | | 313.2 | 134.2 | 447.4 |
| A | RF | Yanco | 2016 | CW | 33.0 | 32.4 | 69.0 | 259.1 | 551.8 | | 551.8 | 174.8 | 726.6 |
| B | RF + WS | Yanco | 2016 | CW | 33.0 | 32.4 | 69.0 | 259.1 | 551.8 | 0 | 551.8 | 174.8 | 726.6 |
| A | RF | Birchip | 2017 | VGB | 26.9 | 28.2 | 35.0 | 221.9 | 280.0 | | 280.0 | 82.6 | 362.6 |
| B | RF + WS | Condoblin | 2017 | CW | 35.4 | 32.3 | 28.3 | 258.2 | 226.4 | 100.0 | 326.4 | 301.8 | 628.2 |
| A | RF | Narrabri | 2017 | HRZ | 46.1 | 36.6 | 29.7 | 293.2 | 237.2 | 20.0 | 257.2 | 356.2 | 613.4 |
| B | RF + WS | Narrabri | 2017 | HRZ | 46.1 | 36.6 | 29.7 | 293.2 | 237.2 | 110.0 | 347.2 | 356.2 | 703.4 |
| A | RF | Yanco | 2017 | CW | 33.0 | 32.4 | 19.8 | 259.1 | 158.6 | | 158.6 | 144.8 | 303.4 |
| B | RF + WS | Yanco | 2017 | CW | 33.0 | 32.4 | 19.8 | 259.1 | 158.6 | X | 158.6 | 144.8 | 303.4 |
| A | RF | Birchip | 2018 | VGB | 26.9 | 28.2 | 14.4 | 221.9 | 115.0 | | 115.0 | 70.8 | 185.8 |
| B | RF + WS | Condoblin | 2018 | CW | 35.4 | 32.3 | 21.8 | 258.2 | 174.3 | 100.0 | 274.3 | 172.9 | 447.2 |
| B | RF + WS | Narrabri | 2018 | HRZ | 46.1 | 36.6 | 26.8 | 293.2 | 214.4 | 148.4 | 362.8 | 203.6 | 566.4 |
| A | RF | Yanco | 2018 | CW | 33.0 | 32.4 | 18.3 | 259.1 | 146.4 | | 146.4 | 141.8 | 288.2 |
| A | RF | Narrabri | 2019 | HRZ | 46.1 | 36.6 | 13.1 | 293.2 | 104.6 | 141.0 | 245.6 | 209 | 454.6 |
| B | RF + WS | Narrabri | 2019 | HRZ | 46.1 | 36.6 | 13.1 | 293.2 | 104.6 | 205.0 | 309.6 | 209 | 518.6 |

Table S8: Yield, Biomass and Water Productivity Best Linear Unbiased Predictions (BLUPs). Values are from 15 trials in four locations. SAL2-type dilutions have resulted in an increase in yields compared to the control Chara variety, while the SAL1-type dilutions have resulted in reduced yields. Values are presented as predictions of random effect of genotype $pm$ standard error. Asterisks indicate significant differences from Chara: * $p < 0.05$, ** $p < 0.01$, *** $p < 0.001$.

| | Trait Measurements | | |
|---|---|---|---|
| Genotype | Yield (T/Ha) | Biomass (T/Ha) | Water Productivity (kg/Ha/mm Water) |
| Chara | 4.70 ± 0.03 | 9.41 ± 0.11 | 14.20 ± 0.10 |
| SAL2 4A.2 | 4.89 ± 0.03 *** | 9.38 ± 0.11 | 14.34 ± 0.10 *** |
| SAL2 4A.3 | 5.09 ± 0.20 | 9.83 ± 0.46 | 14.29 ± 0.51 |
| SAL1 5D.1 | 4.46 ± 0.03 *** | 9.69 ± 0.11 | 13.28 ± 0.10 *** |
| SAL1 5D.2 | 4.57 ± 0.03 *** | 9.78 ± 0.11 * | 13.83 ± 0.10 *** |
| 5D.1x4A.1 | 3.85 ± 0.03 *** | 8.02 ± 0.14 *** | 11.68 ± 0.10 *** |
| 4A.3x5D.1 | 4.31 ± 0.21 | 9.21 ± 0.45 | 13.14 ± 0.46 * |
| 4A.3x5D.1 | 4.39 ± 0.23 | 9.08 ± 0.53 | 13.64 ± 0.99 |
| 5D.1x4A.2 | 4.64 ± 0.21 | 9.39 ± 0.46 | 14.02 ± 0.45 |
| 4A.2x5D.1 | 4.33 ± 0.23 | 9.10 ± 0.53 | 13.71 ± 0.98 |
| 5D.2x4A.3 | 4.94 ± 0.20 | 8.99 ± 0.45 | 13.96 ± 0.46 |
| 5D.2x4A.2 | 4.77 ± 0.20 | 9.35 ± 0.45 | 13.71 ± 0.45 |

### Methods

#### Supporting Method S1 Phylogenetic analysis

Data acquisition:

A list of homologous sequences was retrieved as accessions using the Uniprot web-based BLAST search against the AtSAL1 (NP_201203.3) sequence with the following parameters: blastp, e-value threshold 0.1, restricted to Liliopsida. The full sequences were retrieved from Uniprot (Bateman et al. 2023) using the REST api access. Taxonomic information was retrieved from NCBI Entrez using the Biopython package (Cock et al. 2009). Duplicate sequences and sequences with ambiguous residues were filtered out using a customer python script. AtSAL1 and other paralogous *A. thaliana* reference sequences were added to the data set.

Dataset processing:

An all-v-all BLAST network was built using BLAST+ (NCBI-blast-2.12.0+) (Camacho et al. 2009) with an E-value threshold of 10-10. The network was visualised as a sequence similarity network (SSN) (Atkinson et al. 2009) in Cytoscape 3.9.1 (Shannon et al. 2003). The SSN was filtered by percentage identity edges, removing those with values less than 40%, until the core cluster identified by the *A. thaliana* SAL1/2/3/4 reference sequences was separated from clusters containing the paralogous reference *A. thaliana* reference sequences. Any clusters not connected to the core network were removed.

Sequences making up the core network were filtered to those with amino acid sequence lengths between 250 and 600 aa; the *A. thaliana* reference sequences were then removed. The existing wheat sequences were removed and replaced with their homologues identified from Gramene (Tello-Ruiz et al. 2022). A randomised set of 11 sequences from the SSN cluster from the closely related paralogous AHL enzyme were added to the dataset as an outgroup for the phylogenetic reconstruction.

Sequence alignment:

The sequences were aligned using T-COFFEE Version_13.45.61.3c310a9 (Notredame et al. 2000) and default parameters. The multiple sequence alignment was visualised in Geneious Prime version 2022.0.2. The regions requiring further alignment were realigned using the Geneious implementation of MAFFT version 7.490 (Katoh and Standley 2013) with the algorithm automatically determined, using the BLOSUM80 scoring matrix, a gap penalty of 1.7 and offset value of 0.123, as well as manual curation. Sequences with large insertions or deletions inconsistent with the consensus were removed as possible splice variants.

Phylogenetics: The evolutionary model for phylogenetic reconstruction was found using IQTREE2's ModelFinder (Kalyaanamoorthy et al. 2017), searching against all available protein models. IQTREE2 (Minh et al. 2020) was used for phylogenetic reconstruction, with 10 replicates of each of the two highest scoring evolutionary models from ModelFinder, and each run with 1000 ultrafast bootstraps (Hoang et al. 2018). The best phylogeny was picked by manual evaluation, first removing all trees with a non-contiguous outgroup. The phylogeny was visualised and formatted as a cladogram using the R packages ggtrees (Yu et al. 2017) and ape (Paradis and Schliep 2019).

#### Supporting Method S2 PAP extraction, preparation, quantification by HPLC and SAL activity assay

##### PAP quantification

30-50 mg liquid nitrogen-frozen leaf tissue was ground using a Reich tissuelyser (Qiagen, Hilden, Germany) for 3 min at 25 Hz, using -80°C pre-frozen blocks. Then, 600 $\mu$L of chloroform:methanol (2:1, v:v) and 300 $\mu$L CP buffer (62 mM citric acid, 76 mM $K_2HPO_4$, pH 4) were added and ground for 3 mins at 25 Hz, separately. The homogenate was centrifuged at 17,900 $\times$ g for 10 min at 4°C, and 250 $\mu$L of upper phase was transferred to a new tube and lyophilised overnight. To recover the metabolites, 250 $\mu$L dilute CP buffer (2:3 CP buffer:Water (v:v)) was added and mixed. Fresh weight was converted to dry weight using the calculated relative water content of the sample.

##### SAL activity assay

Protein extraction for SAL activity measurement was performed using 10 mg frozen leaf tissue and ground using the same method as for PAP. Sample extraction buffer (2.5 mM Tris-HCl, pH 7.5, 0.06 mM $\beta$-mercaptoethanol, 1% PVP 360 and 2.5$\times$ Roche proteinase inhibitor cocktail solution) was added and mixed with tissuelyser for 2 min at 30 Hz in a cooled block. Supernatant was recovered and protein quantified by Bradford assay (Bio-Rad Protein Assay Dye Reagent). Proteins were kept on ice and immediately used for activity assay.

SAL activity against PAP was assayed by incubating 0.6 $\mu$g (drought) or 4 $\mu$g (control) protein equivalent from extraction in 10 $\mu$L, with 15 $\mu$L 26.8 $\mu$M PAP standard plus equal volume of 2$\times$ SAL activity buffer (100 mM Tris-MES, pH 7.5, 1 mM $Mg(CH_3COO)_2$). Final reaction volume was 40 $\mu$L. All reactions were prepared on ice. Reaction incubated at 37°C (incubator oven) for 30 min. Reaction stopped by adding 154 $\mu$L CP buffer.

##### HPLC analysis

Prior to HPLC quantification of PAP, 230 $\mu$L of extract was mixed with 20 $\mu$L 45% chloracetaldehyde and incubated for 10 min at 80°C. For assay of SAL activity, the reaction solution was mixed with 10 $\mu$L 45% chloracetaldehyde and incubated for 10 min at 80°C. The derivatisation was stopped by cooling the samples on ice for 15 min. Subsequently, the samples were centrifuged at 17,900 $\times$ g for 10 min at 4°C. The supernatant was transferred to HPLC vials. The commercial PAP (Sigma-Aldrich, A5763) was used as standard for PAP quantification and substrate for SAL activity.

Separation and analysis of the nucleotide derivatives was performed using a reversed-phase HPLC on Kinetex 2.6 $\mu$m XB-C18 100A column (Phenomenex) connected to Agilent 1260 infinity HPLC system and 1100 series fluorescent detector. The gradient for separation of PAP was optimised as follows: column equilibration for 0.05 min with 95% (v/v) of buffer A (5.7 mM Tetrabutylammonium hydrogensulfate (TBAS), pH 5.8) and 5% (v/v) buffer B (67% [v/v] acetonitrile and 33% [v/v] buffer A), followed by linear gradient for 13.44 min to 50% of buffer B, and re-equilibration for 1.81 min with 95% buffer A and 5% buffer B. The chromatograms were recorded and processed with ChemStation software (Agilent).
